# Screening and evaluation of medicine-food homology herbal combinations for ameliorating chemotherapy-induced ovarian aging under an integrated multi-strategy framework

**DOI:** 10.64898/2026.10.22.755587

**Authors:** Jinmei Zhang, Xia Chen, Yahui Cao, Yaou Liu, Nan Ning, Weijie Bian, Xiangning Yang, Yanlei Guo, Jin Zeng

**Affiliations:** Translational Chinese Medicine Key Laboratory of Sichuan Province, Sichuan Academy of ChineseMedicine Sciences, Chengdu, 610041, China; School of Electrical and Information Engineering, Tianjin University, Tianjin, 300072, China; Peking University Chengdu Academy for Advanced Interdisciplinary Biotechnologies, Chengdu, 610213, China; Chongqing Academy of Chinese Materia Medica, Chongqing, 400065, China

**Keywords:** Deep learning, Pretrained models, Herbal combination discovery, Ovarian aging, Medicine–food homology, Astragalus–ginseng

## Abstract

**Background:** Ovarian aging poses a serious threat to women’s reproductive health. Medicine–food homology (MFH) herbs, owing to their favorable safety profile and suitability for long-term use, have become important interventional resources. However, conventional formula screening relies largely on empirical statistics and lacks mechanistic evidence and a systematic screening strategy.

**Methods:** In this study, we integrated single-cell transcriptomic analysis, human-use evidence mining, UPLC/Q-TOF-MS chemical profiling and pretrained protein–molecular representation models to establish a multi-strategy framework for the discovery of MFH herbal combinations. Single-cell transcriptomic datasets related to ovarian aging from the GEO database were analyzed to identify key pathological processes and candidate targets. High-frequency MFH herbs were selected by mining classical prescriptions and clinical literature. By comparing six protein–molecular pretrained representation combinations, ESM-T5 was identified for compound–target affinity prediction and validated by molecular dynamics simulation and surface plasmon resonance (SPR) assays, ultimately identifying the Astragalus–ginseng (Huangqi–Renshen) herb pair (HXF) as the core candidate. Functional expression validation of key targets was performed, and the efficacy of HXF and the expression of related molecules were verified in a D-gal-induced KGN granulosa cell senescence model and a cyclophosphamide-induced rat model of diminished ovarian function.

**Results:** Single-cell analysis revealed that ovarian aging was characterized by impaired steroidogenesis, enhanced MDK–NCL communication between granulosa cells and immune cells, and extracellular matrix remodeling, with CYP19A1, SERPINE1 and MDK as key targets. By further integrating human-use evidence mining and UPLC/Q-TOF-MS compound annotation, an MFH compound library comprising 712 molecules was established. ESM-T5 prediction, molecular dynamics (MD) simulation and SPR assays confirmed that multiple constituents of Astragalus and ginseng exhibited favorable binding activity toward key targets. At the cellular level, knockdown of MDK or SERPINE1 attenuated the senescence phenotype, whereas knockdown of CYP19A1 aggravated it. In vitro and in vivo experiments demonstrated that Astragalus–ginseng ameliorated granulosa cell senescence and diminished ovarian function in rats, restored CYP19A1-related steroidogenesis, downregulated inflammation– and matrix remodeling-related molecules such as SERPINE1 and IL18, and reduced MDK–NCL colocalization signals.

**Conclusion:** This study established a multi-strategy screening framework integrating disease mechanism decoding, human-use evidence constraints and pretrained model prediction, and identified Astragalus–ginseng as a candidate herb pair for ameliorating ovarian aging. Its effects may be associated with promoting steroidogenesis, suppressing SERPINE1/IL18-related inflammation and matrix remodeling, and modulating MDK–NCL signaling. These findings provide a strategic reference for the mechanism-oriented discovery of MFH herbal combinations.

**Graphical abstract:** This study proposes a workflow for MFH herbal formula discovery driven by disease target evidence, constrained by human-use evidence, screened by pretrained models and evaluated through multi-level experiments; applying it to ovarian aging identified Astragalus–ginseng as the core candidate herb pair. Further molecular simulation, functional expression validation, in vitro cell assays and in vivo animal experiments showed that Astragalus–ginseng ameliorated diminished ovarian function and aging-related phenotypes. This provides a scalable strategic reference for the mechanism-oriented screening of MFH herbal combinations and for the development of intervention strategies against complex diseases.

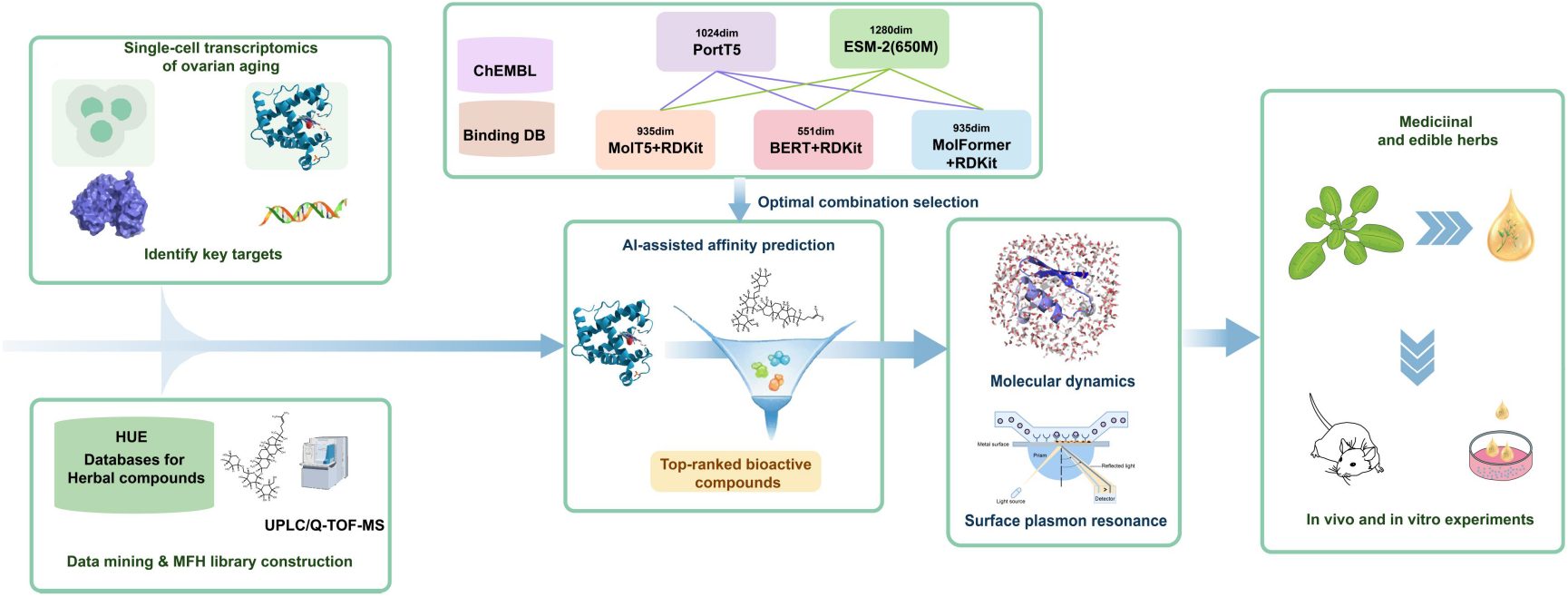

## Introduction

Ovarian aging is characterized by a decline in follicular reserve, functional deterioration of granulosa cells, impaired steroid hormone biosynthesis and an abnormal ovarian microenvironment^[1]^. Ovarian aging can be divided into two categories: physiological aging that occurs naturally with advancing age, and premature ovarian insufficiency and diminished ovarian reserve induced by pathological factors, including genetic disorders, autoimmune diseases, radiotherapy and chemotherapy, which can cause a markedly premature decline in ovarian function^[2]^. It not only leads to reduced fertility and aggravated perimenopausal symptoms, but also increases multi-system health risks such as osteoporosis, metabolic disorders and cardiovascular disease^[3,4]^. Although hormone replacement therapy (HRT) remains the main intervention for alleviating related symptoms, it primarily targets estrogen-deficiency symptoms and has limited effect on improving ovarian reserve and functional decline; controversies regarding long-term breast cancer and cardiovascular risk, together with insufficient patient adherence, further restrict its long-term applicability^[5–8]^. Therefore, exploring alternative strategies with higher safety, suitability for long-term use and action on key steps of ovarian aging is of important clinical and translational significance.

Traditional Chinese medicine (TCM) has accumulated extensive human-use experience in improving diminished ovarian function and regulating women’s reproductive health. Among TCM resources, MFH herbs possess both edible and medicinal attributes with a solid safety foundation, making them important candidate resources for intervening in ovarian aging^[9,10]^. However, MFH medicinal materials are highly diverse, and conventional formula screening relies largely on empirical summarization, usage-frequency statistics and association rules, which makes it difficult to adequately incorporate disease mechanisms. Moreover, TCM is characterized by multi-component, multi-target and multi-pathway actions; relying solely on experience-driven or network association analysis tends to yield an excessively large candidate combination space, homogeneous results and insufficient mechanistic explanation. Therefore, there is an urgent need to establish a systematic screening strategy that integrates disease mechanistic evidence, human-use experience and compound–target prediction capability.

In recent years, public databases represented by the Gene Expression Omnibus (GEO) have accumulated large amounts of disease-related bulk and single-cell transcriptomic data, providing important resources for deciphering cellular heterogeneity, key pathological processes and potential targets^[11]^. Meanwhile, large-scale pretraining approaches such as protein language models and molecular representation models have demonstrated good performance in drug–target interaction prediction, molecular property prediction and virtual screening^[12–16]^. Their combination offers a new route for studying complex TCM systems: single-cell transcriptomics can identify key targets from the disease microenvironment, whereas pretrained models can predict potential active constituents and their targets from TCM compound libraries.

Nevertheless, the implementation of pretrained models in TCM research and development still faces three challenges. First, rebuilding or training large-scale pretrained models places high demands on data, computing power and algorithmic expertise, creating a considerable technical barrier for most pharmaceutical and TCM research settings^[17]^. Second, protein and molecular pretrained models iterate rapidly and differ in representation capability and prediction stability; relying on a single model readily introduces selection bias^[18]^. Third, existing computational tools mostly adopt fixed, platform-based designs, in which built-in models are difficult to replace or extend in a timely manner, limiting the continuous integration and optimization of new models^[19]^. Therefore, how to systematically evaluate and flexibly integrate existing pretrained resources without duplicating large-model development, so as to build an updatable and scalable TCM formula screening framework, remains an unresolved problem.

Therefore, taking ovarian aging as the study scenario, this work constructed an MFH herbal combination screening framework that runs through disease mechanism decoding, human-use evidence constraint, pretrained model screening and multi-level experimental evaluation. Single-cell transcriptomic data were used to decode key pathological processes and identify potential intervention targets; virtual knockout and transient knockdown of key targets were performed to verify their effects on the senescence phenotype; human-use prescription evidence mining for MFH herbs and UPLC/Q-TOF-MS compound identification were integrated to construct an MFH molecular library; different protein and small-molecule representation combinations were compared under a unified evaluation framework to select a better compound–target affinity prediction model for core herb-pair discovery; and the efficacy of candidate herb pairs and the expression of key molecules were validated by molecular dynamics simulation, surface plasmon resonance assays, HPLC-ELSD quantification of constituent contents, and in vitro and in vivo ovarian aging models. This study aims to provide a scalable and updatable methodological reference for the mechanism-oriented screening of MFH herbal combinations in the context of complex diseases.

## 1. Materials and Methods

### 1.1 Animals and cells

Specific pathogen-free (SPF) female Sprague-Dawley rats aged 8 weeks were purchased from Beijing Huafukang Bioscience Co., Ltd. (Beijing, China) and housed in the SPF animal experiment center of the Sichuan Academy of Chinese Medicine Sciences at 22 °C under a 12 h/12 h light/dark cycle with free access to food and water. KGN human ovarian granulosa cells were purchased from Shanghai Fuheng Biotechnology Co., Ltd. (Shanghai, China; FH1125) and cultured in DMEM/F-12 basal medium (Gibco, C11330500BT) supplemented with 10% fetal bovine serum (Viva Cell BIOSCIENCES, C04001-500) and 1% penicillin/streptomycin solution (Gibco, 15140-122) at 37 °C in a 5% CO₂ atmosphere.

### 1.2 Reagents

Biosharp: D-galactose (BS917) and CCK-8 assay kit (BS350B); Beyotime (Shanghai): SA-β-gal staining kit (C0602) and γ-H2AX immunofluorescence staining kit (C2035S); Gibco: Lipofectamine RNAiMAX transfection reagent and Opti-MEM. Sichuan New Lotus Chinese Medicine Decoction Pieces Co., Ltd.: Astragali Radix (2408181), Ginseng Radix et Rhizoma slices (2407029), Rehmanniae Radix Praeparata (2408155), Angelicae Sinensis Radix (D2402019), Glycyrrhizae Radix et Rhizoma slices (2404004), Poria (2408130), Dioscoreae Rhizoma (2409072), Corni Fructus (2406194), Lycii Fructus (2408008) and Rehmanniae Radix (2509100); MedChemExpress (MCE, USA): astragaloside IV (HY-N0431), isoastragaloside II (HY-N0888), ginsenoside Rd (HY-N0043) and ginsenoside Re (HY-N0044) reference standards. Chromatographic columns: Kinetex XB-C18 (100 mm × 2.1 mm, 2.6 μm, Phenomenex, USA) and PY-C18 (4.6 mm × 250 mm, 5 μm). Recombinant proteins for SPR: CYP19A1 (PA2000-4059, Hubei Aiputi Biotechnology), MDK (10247-hnab, Beijing Sino Biological) and PAI-1 (rp181294, Aladdin); the CM5 sensor chip (29-1049-88) was purchased from Cytiva (USA). Chengdu Rongwei Gene: RNA extraction (K101-01), reverse transcription (A504-02) and PCR (A403-01) kits; total RNA from ovarian tissue was extracted using TRK Lysis Buffer. Cyclophosphamide for injection (4B660A, Baxter Oncology GmbH, Germany), Progynova (869A, Bayer HealthCare) and isoflurane (2024052601, Shandong Ante Animal Science and Technology Co., Ltd.). Antibodies: Immunoway (USA) — CYP19A1 (YT1190), KI67 (YM8189), γ-H2AX (YM8195), p-CREB (YM8632), FN1 (YM8309), IL-1β (YT5201) and MDK (YM960025); Abcam (Cambridge, UK) — AMH (Ab313767), NCL (ab129200) and PAI-1 (ab222754); Huabio — IL-18 (HA723978); Cell Signaling Technology (CST) — HRP-conjugated goat anti-rabbit secondary antibody (7074P2); Servicebio — FITC (GB22303)– and CY3 (GB21303)-conjugated goat anti-rabbit secondary antibodies and antifade mounting medium. Elabscience (USA): ELISA kits for serum FSH (E-EL-R0391), LH (E-EL-R0026), AMH (E-EL-R3022) and E2 (E-OSEL-R0001).

### 1.3 Instruments

Thermo Fisher Scientific: 311 water-jacketed CO₂ incubator, 1820 microplate reader, 2720 Thermal Cycler (RT-qPCR), NX50 cryostat, A80400011 antigen retrieval instrument and Qubit 3.0; Leica: 090-135008 inverted microscope (Leica, Germany) and RM2016 microtome (Leica, Shanghai). Shimadzu (Japan): LC30A ultra-high performance liquid chromatograph; AB SCIEX (USA): TripleTOF 4600 quadrupole–time-of-flight high-resolution mass spectrometer; Waters (USA): 2695 high performance liquid chromatograph; Cytiva (USA): Biacore 8K surface plasmon resonance system. SCILOGEX: SCI1000-G gradient PCR instrument; Shanghai Tanon: 4600 chemiluminescence imaging system. Changzhou Zhongwei: TSJ-II tissue dehydrator, PHY-III tissue floatation and drying station and BMJ-III embedding machine; Motic: BA410 microscope. Agilent: Agilent 2100 Bioanalyzer; MGI: DNBSEQ-T7 sequencing platform (PE150).

## 2. Methods

### 2.1 Single-cell transcriptomic integration analysis

Human ovarian aging single-cell transcriptomic datasets (GSE255690, GSE202601) were obtained from the GEO database (https://www.ncbi.nlm.nih.gov/geo), including ovarian samples from young women (18–29 years, n = 7) and aged women (47–54 years, n = 7). Use of the public data complied with the GEO human subjects protection policy and required no additional ethical approval. Analysis was performed using R (v4.3.2) and Seurat: Seurat objects were constructed (min.cells = 3, min.features = 300), 2,000 highly variable genes were selected after LogNormalize normalization, batch effects were corrected using a CCA anchor-based integration strategy, and cell cycle effects were regressed out using CellCycleScoring and ScaleData. PCA, K-nearest neighbor graph construction, unsupervised clustering and UMAP visualization were performed on the integrated data, with final parameters set to dims = 35 and resolution = 0.2. FindAllMarkers was used to identify cluster marker genes (Wilcoxon test, logFC > 0.25, adjusted P < 0.05), and cell type annotation was completed by combining CellMarker 2.0 (http://117.50.127.228/CellMarker/), the original publications and canonical marker genes^[20,21]^. FindMarkers was used to screen differentially expressed genes between the young and old groups (min.pct = 0.2, |log2FC| ≥ 1, adjusted P < 0.05), and GO/KEGG enrichment, STRING protein–protein interaction network and Cytoscape topological analyses were conducted^[22,23]^. Candidate genes were scored comprehensively based on expression difference, statistical significance, proportion of expressing cells, senescence gene databases and evidence related to ovarian function. CellChat was used to construct and compare cell–cell communication networks between the two groups based on CellChatDB.human (excluding cell populations with fewer than 10 cells), to identify differential communication pathways and key cell interactions; differential expression, enrichment and network analyses were performed on cell subsets with marked communication changes to screen subset-specific key genes.

### 2.2 Virtual knockout

To assess the reliability of SERPINE1, MDK and NCL as potential targets of ovarian aging, virtual knockout analysis was performed using scTenifoldKnk^[24]^. The granulosa cell subset was extracted, 5,000 highly variable genes were selected, the cell number was set to 2,500, and scTenifoldKnk was run to obtain the differentially regulated results before and after virtual knockout.

### 2.3 Cell culture and modeling

KGN human ovarian granulosa cells (Shanghai Fuheng Biotechnology Co., Ltd., FH1125) were cultured in DMEM/F-12 supplemented with 10% fetal bovine serum and 1% penicillin/streptomycin at 37 °C in a 5% CO₂ atmosphere. KGN cells were treated with D-gal (72 mg/mL) for 24 h to induce cellular senescence^[25]^.

### 2.4 siRNA transient transfection

KGN cells were seeded in 6-well plates at 2 × 10⁵ cells per well and cultured in antibiotic-free medium until reaching approximately 70% confluence before transfection. Two independent siRNAs were designed for each of MDK, SERPINE1 and CYP19A1, and a non-targeting siRNA (si-NC) was used as a negative control (Tab. 1). Transient transfection was performed using Lipofectamine RNAiMAX: siRNA and transfection reagent were separately diluted in Opti-MEM, incubated at room temperature for 30 min to form complexes, and then added to the cell culture medium at a final siRNA concentration of 50 nM. At 48 h after transfection, RT-qPCR and Western blot were used to detect target gene mRNA and protein expression to evaluate knockdown efficiency.

**Table. 1.**
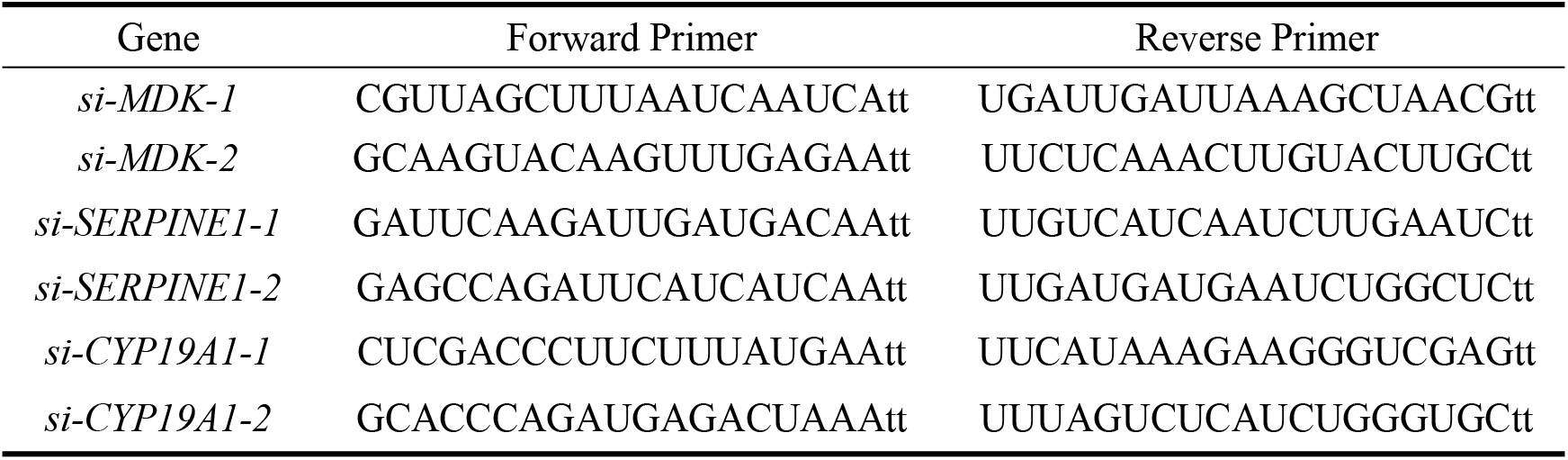
siRNA Sequences.

### 2.5 Evaluation of cellular senescence phenotype after gene knockdown

At 48 h after transfection, cellular senescence was induced. A normal control group, an si-NC group, a D-gal model group and D-gal combined with each target gene siRNA transfection group were set up. After treatment, cell viability was measured using a CCK-8 kit. The degree of cellular senescence was assessed using an SA-β-gal staining kit, and the proportion of SA-β-gal-positive cells was quantified using ImageJ^[25]^.

### 2.6 Data mining to identify MFH herbs for ameliorating ovarian aging

CNKI (https://www.cnki.net/), Wanfang (https://www.wanfangdata.com.cn/), VIP (https://qikan.cqvip.com/) and SinoMed (https://www.sinomed.ac.cn/zh/) databases were systematically searched from database inception to February 2025, using search terms such as ovarian aging/ovarian ageing/premature ovarian insufficiency and Chinese herbal compound/formula/anti-aging, and candidate herbs were screened according to the medicine–food homology catalogue of the National Health Commission. Clinical studies and classical formula literature using oral Chinese herbal compounds to treat ovarian aging or premature ovarian insufficiency, with clearly defined prescription composition and evaluable efficacy, were included; single herbs, incomplete or duplicated prescriptions, reviews and non-clinical studies were excluded. Herb names were standardized according to the Chinese Pharmacopoeia^[26]^. Herb usage frequency (≥ 20), properties and flavors, meridian tropism and efficacy were analyzed^[27]^; association rule analysis and hierarchical clustering were performed using R (v4.3.2), with support ≥ 10% and confidence ≥ 60%. Based on herb frequency, compatibility association and clustering results, the top 10 key MFH herbs for ameliorating ovarian aging were selected.

### 2.7 Preparation of herbal decoctions

Each herbal decoction piece (10.0 g) was soaked in 10 volumes of water for 30 min and then decocted twice: the first decoction was simmered for 60 min after boiling, and the second was simmered for 30 min after adding another 10 volumes of water. The filtrates were combined, concentrated by rotary evaporation, adjusted to a final volume of 100 mL, aliquoted, and stored at −80 ℃.

### 2.8 UPLC/Q-TOF-MS chemical profiling

A 200 µL aliquot of the decoction was mixed with 400 µL methanol, vortexed for 30 s and centrifuged at 12,000 rpm for 15 min, and the supernatant was subjected to UPLC/Q-TOF-MS analysis with an injection volume of 2 µL. A Kinetex XB-C18 column (100 × 2.1 mm, 2.6 µm) was used with a mobile phase of 0.1% formic acid in water (A) and acetonitrile (B) at a flow rate of 0.25 mL/min and a column temperature of 30 °C. The ESI gradient elution program was: 0–1 min, 5% B; 1–7 min, 5%–40% B; 7–11 min, 40%–55% B; 11–21 min, 55%–85% B; 21.01–25 min, 5% B. The APCI gradient was: 0–1 min, 55% B; 1–4 min, 55%–90% B; 4–7 min, 90% B; 7–7.5 min, 90%–55% B; 7.5–10 min, 55% B. Data were acquired on a TripleTOF 4600 mass spectrometer in both positive and negative ion modes over a scan range of m/z 100–1000; MS/MS fragmentation spectra were obtained by information-dependent acquisition (IDA) for compound identification.

Candidate reference compounds were obtained from the HERB (http://herb.ac.cn/) and TCMSP (https://www.tcmsp-e.com) databases and literature searches. Raw data were processed using PeakView 1.2, and preliminary screening was based on accurate mass error |ppm| ≤ 15, retention time, isotope distribution and MS/MS fragmentation information; compounds were further confirmed by combining the MassBank spectral library and fragmentation rules, thereby constructing a traceable MFH compound molecular library for subsequent affinity prediction and experimental validation.

### 2.9 Modular integration and evaluation of pretrained models

Molecular embeddings were generated using ChemBERTa ^[28]^, MolFormer^[29]^ and MolT5^[30]^, respectively, and RDKit was used to calculate physicochemical and topological descriptors of the molecules, which were concatenated with the molecular embeddings to supplement interpretable physicochemical prior information; protein representations were generated using ESM-2 650M^[31]^and ProtT5^[32]^ to produce sequence embeddings of target proteins. The three molecular representations and two protein representations were fully combined to form six protein–molecular representation configurations. For each configuration, protein and molecular embeddings were concatenated and fed into the same downstream regression architecture to predict compound–target binding affinity.

Model training data were obtained from the ChEMBL and BindingDB databases^[33]^. Molecule–protein interaction data with experimentally determined KD values were collected; after molecular structure standardization, protein sequence matching, and removal of duplicates and outliers, 149,994 valid molecule–protein affinity samples were obtained. The dataset was randomly split into training, validation and test sets at a ratio of 8:1:1. To evaluate model robustness, five-fold cross-validation was introduced in the training data to examine model performance under different data partitions. Before model training, KD values were uniformly converted to pKD values, i.e., pKD = −log10(KD, mol/L). Model performance was comprehensively evaluated using the coefficient of determination, mean absolute error, mean squared error and root mean squared error, denoted as R², MAE, MSE and RMSE, respectively^[34]^. The representation combination with better overall performance was used for subsequent compound–target affinity prediction on the MFH molecular library. All training and prediction tasks were completed on an Intel Xeon E5-2686 v4 CPU and an NVIDIA Tesla P100 GPU.

**Fig. 1.**
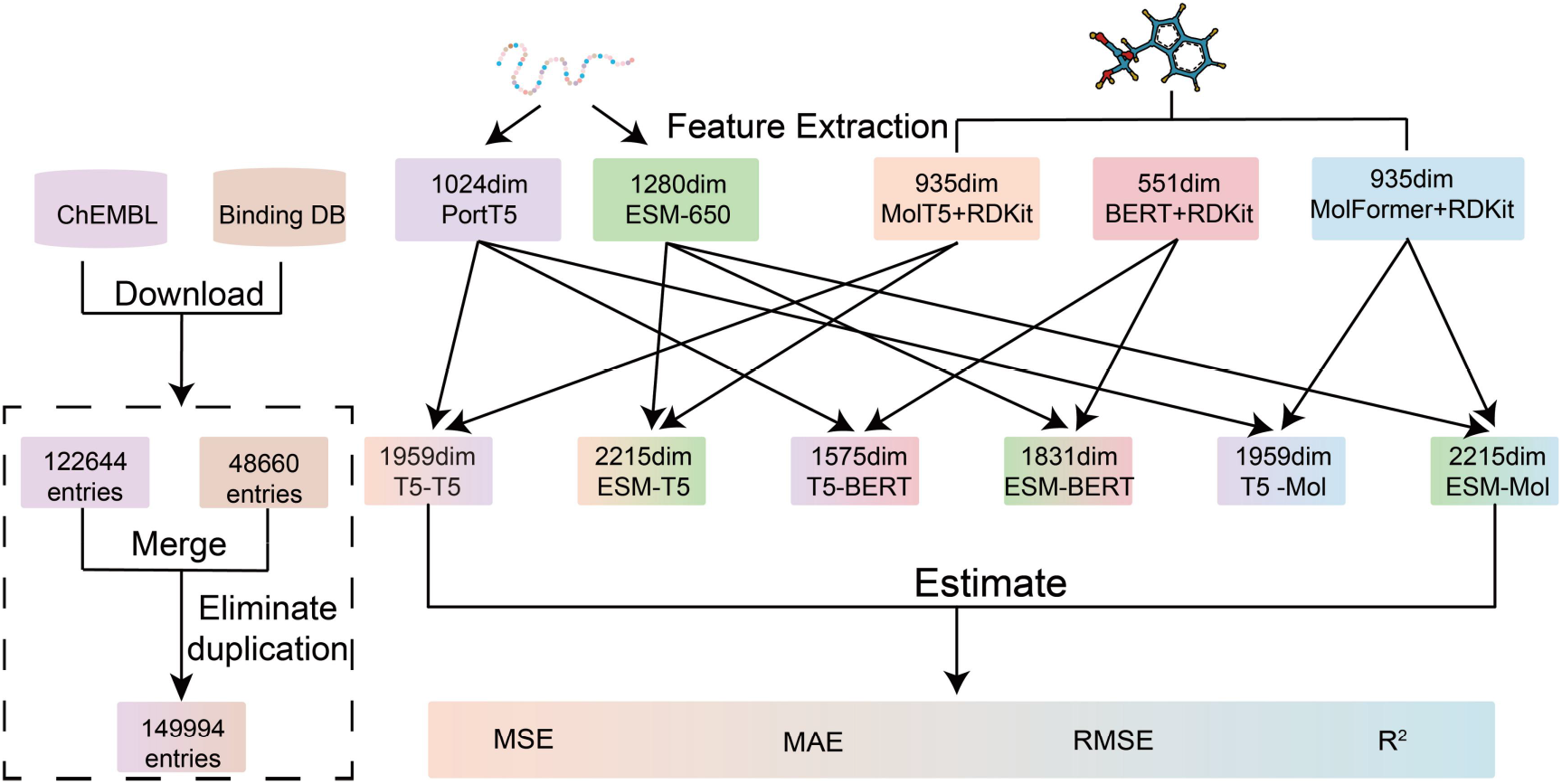
Modular integration and evaluation of pretrained protein and molecular representations for affinity prediction.

**Fig. 1.**
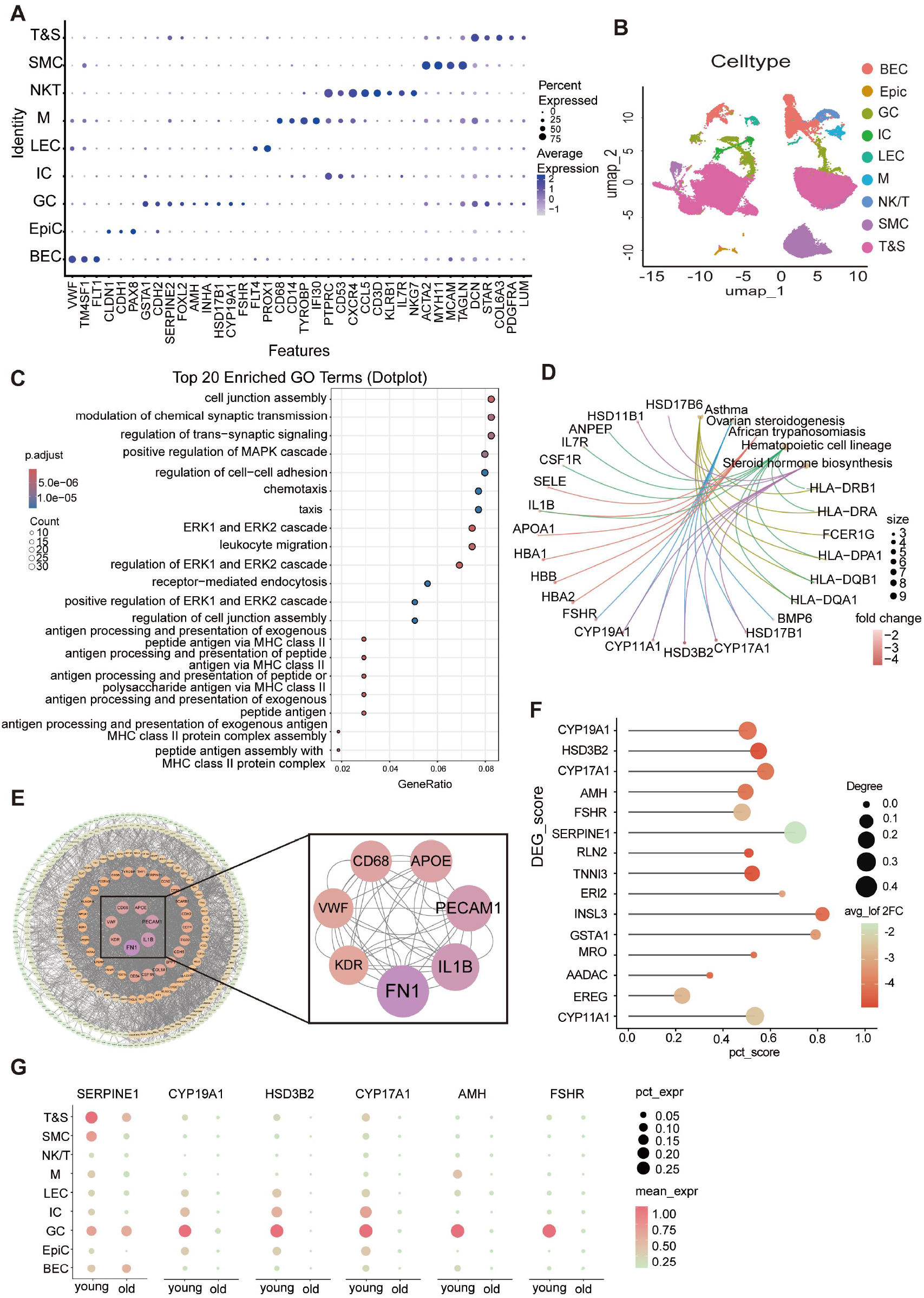
Single-cell transcriptomic analysis identifies ovarian aging-associated transcriptional alterations and candidate regulatory nodes. (A) UMAP visualization of ovarian cells from young and old groups after quality control, normalization and batch correction. (B) Annotation of major ovarian cell populations, including blood endothelial cells (BECs), epithelial cells (EpiCs), granulosa cells (GCs), immune cells (ICs), lymphatic endothelial cells (LECs), macrophages (Ms), NK/T cells, smooth muscle cells (SMCs), and theca and stromal cells (T&S cells). (C) Differentially expressed genes (DEGs) between young and old ovarian cells. (D) Functional enrichment analysis of aging-associated DEGs, highlighting biological processes related to leukocyte migration and chemotaxis, cell junction and adhesion, ovarian steroidogenesis, steroid hormone biosynthesis, MAPK/ERK signaling, and MHC II-associated antigen processing and presentation. (E) Protein–protein interaction network analysis of aging-associated DEGs, showing FN1 and IL1B as highly connected nodes. (F) Integrated prioritization of candidate aging-associated genes based on logFC, statistical significance, aging-related annotation, PubMed evidence, and network topology. (G) In silico perturbation analysis of SERPINE1 in granulosa cells, identifying markedly affected downstream genes such as MAP3K5 and DOCK5. (H) Cell type-specific expression and detection rate of SERPINE1 in young and old groups, showing increased SERPINE1 expression in blood endothelial cells and granulosa cells from aged ovaries.

### 2.10 Compound–target protein affinity prediction

Compounds from the 10 key herbs were deduplicated, structurally standardized and converted into SMILES format to construct an MFH molecular library for RDKit descriptor calculation and affinity prediction. Compounds in the library were paired one by one with candidate target proteins, and the optimized protein–ligand affinity prediction model was used to calculate the predicted binding affinity (pK_D) for each compound–target combination.

### 2.11 Molecular dynamics simulation

Molecular dynamics (MD) simulations of candidate protein–ligand complexes were performed for 100 ns using GROMACS 2020.3. The protein and ligand were parameterized with the AMBER ff14SB and GAFF2 force fields, respectively. The system was built in a truncated octahedral TIP3P water box with a boundary distance of 1.0 nm, and Na⁺/Cl⁻ were added to achieve charge neutralization. After energy minimization, 200 ps of heating, 500 ps of NVT equilibration and 500 ps of NPT equilibration, a 100 ns production simulation was performed, with trajectories saved every 10 ps. Stable trajectories from the 90–100 ns interval were selected, and the MM/GBSA method was used to calculate protein–ligand binding free energies and their energy components.

### 2.12 Surface plasmon resonance (SPR) assay

Experiments were performed at 25 °C. Recombinant proteins at a concentration of 20 ng/µL were immobilized on the surface of a CM5 sensor chip by amine coupling. Test small molecules were prepared in a series of concentrations in running buffer containing 5% DMSO and sequentially injected over the chip, and real-time association and dissociation signals were recorded. After reference flow cell subtraction and blank correction, sensorgrams were fitted using Biacore Insight Evaluation Software and the equilibrium dissociation constant (K_D_) was calculated.

### 2.13 Core herb pair screening

The top 10 differentially expressed genes ranked by global priority, the PPI network core targets FN1 and IL1B, and the key ligand–receptor pair MDK–NCL were selected as candidate targets; for each target, the top 10 candidate active compounds were screened according to predicted pK_D values from high to low, and the compound–target affinity profile was displayed as a heatmap. Based on the screened high-affinity compounds and their source herbs, a three-layer herb–compound–target network was constructed and visualized using Cytoscape 3.10.1. Considering network coverage together with the enrichment of high-affinity molecules at the target level, the core herb pair for ameliorating ovarian aging, namely Astragalus and ginseng (HXF), was identified.

### 2.14 Quantification of four saponins in HXF by HPLC-ELSD

The HXF decoction was prepared from Astragalus and ginseng at a mass ratio of 1:1^[26,35]^ using the same procedure as described in section 1.7. High performance liquid chromatography with evaporative light scattering detection (HPLC-ELSD) was used for the simultaneous determination of ginsenoside Re, astragaloside IV, ginsenoside Rd and isoastragaloside II. The chromatographic system was a Waters 2695 HPLC instrument equipped with a PY-C18 column (4.6 mm × 250 mm, 5 μm), with a column temperature of 35 °C, a flow rate of 1.0 mL/min and an injection volume of 20 μL. Gradient elution was performed with acetonitrile (A) and 0.2% formic acid aqueous solution (B): 0–10 min, 18%–20% A; 10–30 min, 20%–23% A; 30–38 min, 23%–35% A; 38–50 min, 35% A; 50–65 min, 35%–90% A; 65–75 min, 90% A, followed by re-equilibration to the initial conditions. The ELSD gas flow rate was 2.7 L/min and the drift tube temperature was 100 °C. Three batches of HXF decoction (HXF1, HXF2, HXF3) were centrifuged at 12,000 r/min for 20 min, and the supernatants were filtered before injection. Ginsenoside Re (20.41 mg), astragaloside IV (2.43 mg), ginsenoside Rd (2.15 mg) and isoastragaloside II (1.15 mg) were accurately weighed, dissolved in methanol to a final volume of 5 mL as a mixed stock solution, and 0.06–0.40 mL aliquots were serially diluted to 2 mL to obtain a series of standard solutions; standard curves were plotted as lg C versus lg A. For method validation, linearity was evaluated by R² ≥ 0.99, precision by the RSD of contents obtained from six consecutive injections of the HXF2 test solution, and LOD and LOQ by injection amounts at signal-to-noise ratios of 3 and 10, respectively.

### 2.15 Cell viability assessed by CCK-8 assay

KGN cells in the logarithmic growth phase were seeded in 96-well plates at 1 × 10⁴ cells per well. Except for the normal control and blank groups, all other groups were treated with DMEM/F-12 complete medium containing D-gal (72 mg/mL) and administered HXF decoction at 10, 20, 40, 80 and 160 µg/mL. After treatment for 24, 48 and 72 h, 10 µL of CCK-8 reagent was added to each well and incubated at 37 °C in the dark for 2 h, and the absorbance (OD) was measured at 450 nm. Based on the cell viability results, the appropriate HXF intervention concentration and treatment duration for subsequent experiments were determined.

### 2.16 Senescence-associated β-galactosidase (SA-β-gal) staining

KGN cells in the logarithmic growth phase were seeded in 6-well plates at 2 × 10⁵ cells per well and cultured for 24 h. After 24 h of senescence induction, cells were treated with HXF decoction (20, 40 and 80 µg/mL) for 24 h. After treatment, cells were washed with PBS and fixed with fixative at room temperature for 15 min, followed by three PBS washes. SA-β-gal staining working solution was then added and incubated at 37 °C in the dark overnight. The staining solution was discarded, cells were washed 2–3 times with PBS, images were acquired under a microscope, and the proportion of SA-β-gal-positive cells was quantified using ImageJ.

### 2.17 γ-H2AX immunofluorescence staining

The treatment groups were the same as those for SA-β-gal staining. Cells were gently washed with PBS and fixed at room temperature for 15 min. After fixation, cells were washed, blocked with immunostaining blocking solution at room temperature for 120 min, and then incubated with rabbit anti-γ-H2AX primary antibody at 4 °C overnight. The next day, after washing, cells were incubated with Alexa Fluor 488-conjugated anti-rabbit secondary antibody at room temperature in the dark for 1 h. Nuclei were stained with DAPI, slides were mounted with antifade mounting medium, and images were acquired under a fluorescence microscope.

### 2.18 Animal modeling and drug administration

Thirty-six rats with regular estrous cycles were selected and randomly divided into groups of six. Rats in the modeling group received a single intraperitoneal injection of cyclophosphamide (CTX, 140 mg/kg) to establish the ovarian aging model, whereas control rats received an equal volume of normal saline. After successful modeling^[36]^, model rats were randomly divided into a model group, a 2.1 g/kg HXF group, a 4.2 g/kg HXF group, an 8.4 g/kg HXF group and a 0.105 mg/kg Progynova group. The low dose of HXF (2.1 g/kg) and the dose of Progynova (0.105 mg/kg) were calculated from the human dose according to the body surface area conversion factor. Each treatment group received intragastric administration once daily for 14 consecutive days. The normal control and model groups were given distilled water by gavage at 1 mL/100 g body weight daily. During the experiment, changes in body weight and estrous cycles were recorded continuously.

### 2.19 Estrous cycle examination and organ indices

Vaginal exfoliated cells were gently collected daily at 09:00 using a sterile cotton swab moistened with normal saline, evenly smeared onto glass slides, dried, stained with methylene blue solution for 10 min, and observed under a light microscope. Estrous cycle stages were determined as follows: proestrus, predominantly nucleated epithelial cells; estrus, predominantly cornified anucleated squamous epithelial cells; metestrus, a mixture of cornified epithelial cells and leukocytes; diestrus, predominantly leukocytes. Rats remaining in the same stage for 5 consecutive days were judged to have estrous cycle disturbance^[37]^.At 24 h after the last administration, rats were anesthetized by intraperitoneal injection of sodium pentobarbital (50 mg/kg), and blood was collected from the abdominal aorta before euthanasia. The ovaries, liver, spleen and kidneys were rapidly isolated, and organ indices were calculated as follows: organ index (mg/g) = organ wet weight (mg)/body weight (g).

### 2.20 Serum hormone measurement

Collected whole blood was allowed to stand for 30 min and then centrifuged at 1,000 g for 15 min to obtain the supernatant. Serum levels of FSH, LH, E2 and AMH were measured strictly according to the ELISA kit instructions. Absorbance was read at 450 nm using a microplate reader, and standard curves were fitted and sample concentrations calculated using ELISACalc software.

### 2.21 Hematoxylin and eosin (H&E) staining

Ovarian tissues were fixed in 4% paraformaldehyde for 24 h, dehydrated through a graded ethanol series, cleared in xylene and embedded in paraffin, and serial sections (3 µm thick) were prepared. After dewaxing and rehydration, sections were stained with hematoxylin and eosin, dehydrated through graded ethanol, cleared in xylene and mounted with neutral resin. Ovarian tissue morphology was observed under a light microscope, and follicles at different developmental stages (primordial, primary and atretic follicles) were classified and counted according to morphological features ^[38]^.

### 2.22 Immunohistochemistry

Paraffin sections were dewaxed and rehydrated, and antigen retrieval was performed using citrate buffer. After cooling to room temperature, sections were treated with 3% hydrogen peroxide to block endogenous peroxidase and with 5% goat serum to block non-specific binding. Corresponding primary antibodies were then added and incubated overnight at 4 °C in a humidified chamber. The next day, after washing with PBS, sections were incubated with HRP-conjugated secondary antibody at room temperature and developed with DAB. Finally, nuclei were counterstained with hematoxylin, and sections were dehydrated, cleared, mounted and imaged under a light microscope.

### 2.23 Immunofluorescence

Paraffin sections were dewaxed, rehydrated and subjected to antigen retrieval, blocked with 5% goat serum for non-specific binding, and then incubated with corresponding primary antibodies. After washing with PBS, sections were incubated with fluorescently labeled secondary antibodies in the dark and nuclei were stained with DAPI. For multiplex immunofluorescence staining, a cyclic staining strategy was used for multi-marker detection. After staining, sections were mounted with antifade mounting medium and imaged under a fluorescence microscope. Staining intensity was quantified using ImageJ.

### 2.24 Transcriptome sequencing

Ovarian tissue (15–30 mg) was ground in liquid nitrogen and total RNA was extracted using TRK Lysis Buffer. After homogenizer column purification, RNA binding column adsorption and washing, RNA was eluted with nuclease-free water. RNA quality was assessed using NanoDrop, Qubit and agarose gel electrophoresis. Libraries were constructed using the VAHTS Universal V6 RNA-seq Library Prep Kit for MGI (Vazyme, NRM605), and after quality control on an Agilent 2100 Bioanalyzer, PE150 sequencing was performed on the DNBSEQ-T7 platform. Low-quality reads, adapter-contaminated reads and short reads were filtered out to obtain clean reads. Gene expression was quantified using RSEM, and sample consistency was evaluated by Pearson correlation analysis and PCA. Differentially expressed genes were identified using DESeq2 with thresholds of |log2FC| ≥ 1 and FDR < 0.05. GO and KEGG functional enrichment analyses were performed using clusterProfiler, with FDR < 0.05 considered significant.

### 2.25 Real-time quantitative polymerase chain reaction (RT-qPCR)

Total RNA was extracted using a cell/tissue total RNA extraction kit. One microgram of total RNA was reverse-transcribed into cDNA using the MetaScript qPCR RT Super Mix with dsDNase kit. Quantitative analysis was performed using the SYBR qPCR Master Mix Kit, with GAPDH as the internal reference gene, and relative expression levels were calculated using the 2^−ΔΔCT method. Primer sequences are listed in Tab. 2.

**Table 2.**
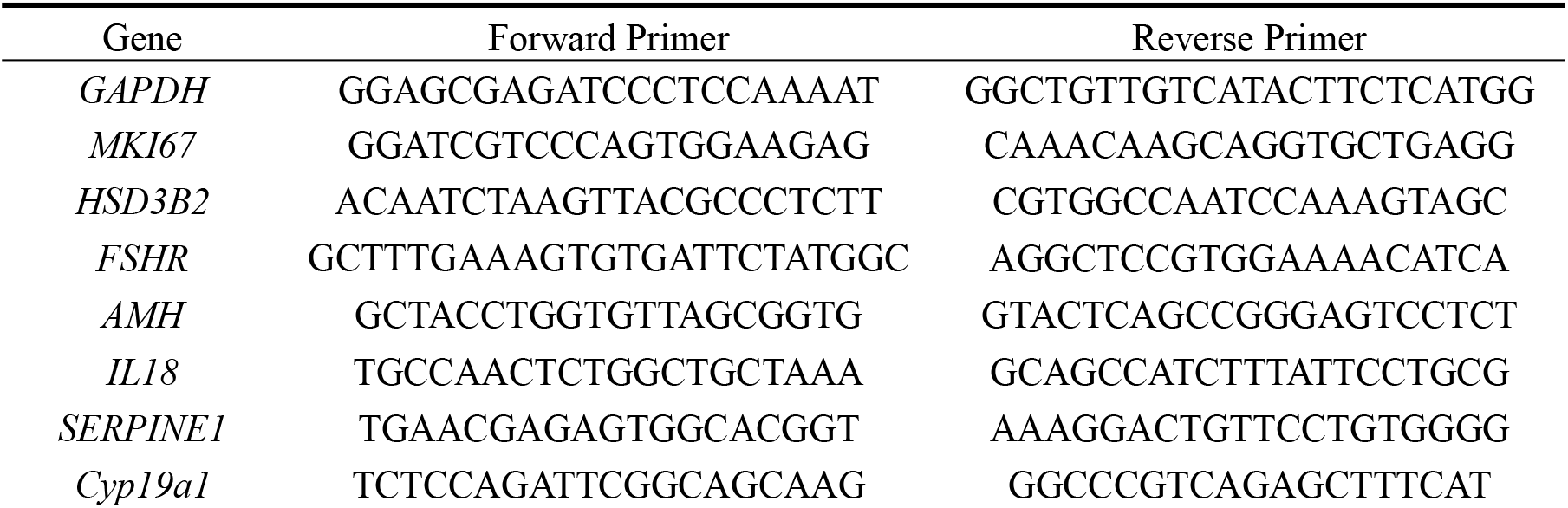
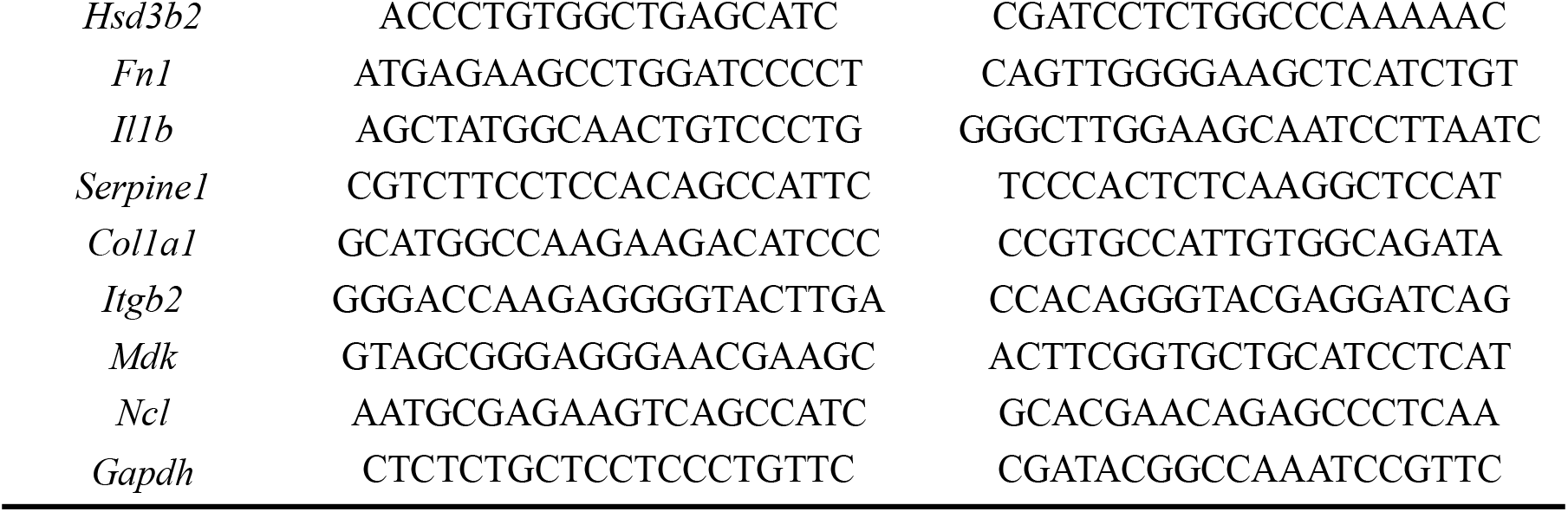
Primers used for PCR.

### 2.26 Western blotting

Ovarian tissue was lysed in RIPA lysis buffer containing protease and phosphatase inhibitors (100:1:1) at a ratio of 0.1 g:1 mL, homogenized three times for 30 s each, lysed on ice for 15 min, and centrifuged at 13,000 rpm for 15 min at 4 °C, and the supernatant was collected. Protein concentration was determined by the BCA method, 1/4 volume of 5× loading buffer was added, and samples were denatured at 100 °C for 10 min. Proteins were separated by 12% SDS-PAGE and transferred to membranes; after rapid blocking, membranes were incubated with primary antibodies at 4 °C overnight and with secondary antibodies at room temperature for 1 h. Signals were developed with ECL, and band gray values were analyzed using ImageJ.

### 2.27 Statistical analysis

Statistical analysis was performed using GraphPad Prism 10.1.2. Values are expressed as mean ± standard deviation (SD). Differences among groups were assessed using one-way analysis of variance, with a significance level of P < 0.05. Data visualization was performed using GraphPad Prism 10.12 and R 4.3.2.

## 3. Result

### 3.1 Single-cell transcriptomics reveals altered cellular composition in ovarian aging and identifies key candidate targets

To decipher the molecular features of ovarian aging at the cellular level, quality control, normalization and batch effect correction were performed on ovarian aging-related scRNA-seq data, and 42,345 cells from the young group and 48,619 cells from the old group were finally included for subsequent analysis (Fig. S1A-D). The first 1–50 principal components were selected by principal component analysis and UMAP dimensionality reduction clustering was performed, identifying 20 cell clusters (Fig. S1E, F); based on marker gene expression, nine major cell populations were finally annotated (Fig. 1A, B). Among them, ICs were a small population identified after data integration that expressed immune cell signature genes but lacked other specific marker genes, and were therefore annotated separately.

On this basis, differential expression between the young and old groups was compared at the global cell level, and 480 significantly differentially expressed genes (DEGs) were identified (Fig. S2). Functional enrichment analysis showed that these DEGs indicated a transcriptional signature of the ovarian aging tissue microenvironment characterized by enhanced immune recruitment, altered cell adhesion and aberrant steroidogenesis (Fig. 1C, D). The PPI network constructed from the DEGs showed that FN1 and IL1B occupied highly connected regions of the network, suggesting that they may participate in the global transcriptional network remodeling associated with ovarian aging (Fig. 1E). Weighted comprehensive scoring of the DEGs showed that top-ranked genes including CYP19A1, HSD3B2, CYP17A1, AMH and FSHR were related to ovarian steroidogenesis and follicular function^[39,40]^, whereas SERPINE1 may represent a candidate regulatory node in the directions of aging-related inflammation, fibrinolytic homeostasis and matrix remodeling^[41]^(Fig. 1F-H).

To decipher the changes in intercellular interactions during ovarian aging, cell–cell communication inference analysis was further performed. The results showed that, compared with the young group, the old group had a reduced overall number of interactions but increased communication strength, with markedly enhanced communication between granulosa cells (GCs) and NK/T cells and macrophages (Ms) (Fig. 2A, B). Ligand–receptor expression analysis revealed that the expression of the MDK–NCL ligand–receptor axis between GCs and multiple cell types including BECs, NK/T cells, Ms and SMCs was significantly elevated in the old group (Fig. 2C), indicating that ovarian aging is accompanied by remodeling of cell communication patterns characterized by aberrant granulosa cell–immune cell interactions, in which the MDK–NCL axis may be a key communication pathway in the ovarian aging microenvironment.

**Fig. 2.**
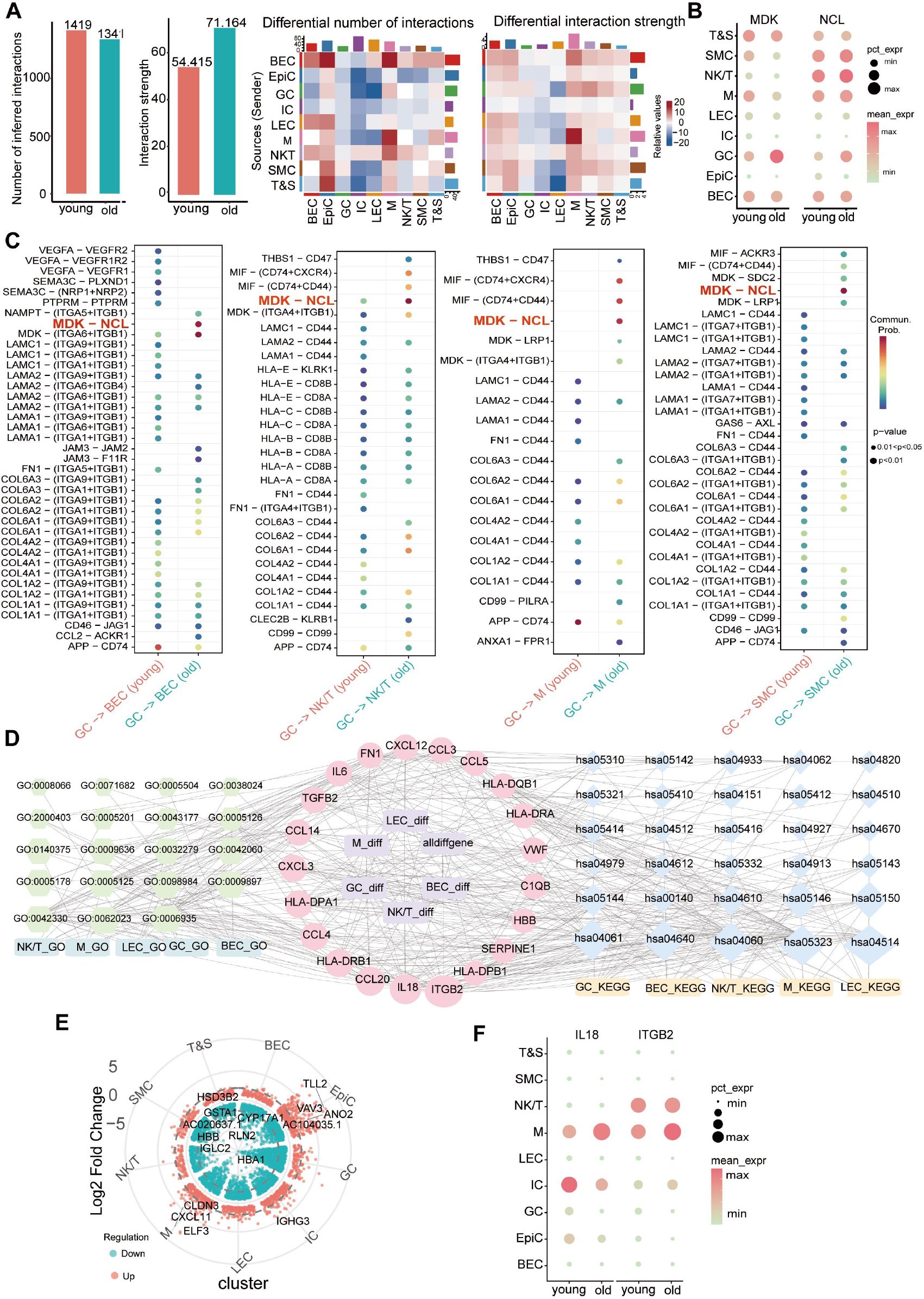
Cell–cell communication analysis reveals enhanced MDK–NCL signaling and immune-related microenvironmental remodeling in aged ovaries. (A) Global comparison of cell–cell communication between young and old groups, showing reduced overall interaction number but increased communication strength in aged ovaries, particularly between granulosa cells and immune cell populations. (B) Ligand–receptor interaction analysis identifying enhanced MDK–NCL signaling between granulosa cells and multiple cell populations in the old group. (C) Visualization of MDK–NCL-mediated intercellular communication patterns in young and old ovaries. (D) Cell type-specific differential expression analysis identifying aging-associated genes in distinct ovarian cell populations. (E) Functional enrichment analysis of cell type-specific aging-associated genes, indicating immune activation, inflammatory signaling and extracellular matrix-related remodeling. (F) Prioritization of cell type-specific candidate nodes, including ITGB2 and IL18.

For the key cell types identified by the above communication analysis, a potential interaction network was constructed based on differentially expressed genes and the top 10 enriched pathways ranked by significance. The results showed that ITGB2 and IL18 were located at the center of the network (Fig. 2D), and both were significantly upregulated in macrophages of the old group, while ITGB2 was also upregulated in NK/T cells (Fig. 2E, F). Taken together, candidate key targets related to ovarian aging were preliminarily determined: CYP19A1, HSD3B2 and FSHR were mainly associated with steroidogenesis and follicular dysfunction; MDK and NCL participated in aberrant cell communication in ovarian aging; ITGB2, IL18 and IL1B reflected immune activation and inflammatory amplification; and SERPINE1 was involved in fibrinolytic homeostasis imbalance, extracellular matrix remodeling and transcriptional network hub regulation. These targets collectively provide the basis for subsequent target functional validation and compound–target prediction.

### 3.2 Virtual knockout and transient knockdown jointly validate the functions of key targets in ovarian aging

To evaluate the regulatory impact of candidate targets at the network level, virtual knockout analysis of granulosa cells was performed using scTenifoldKnk^[42]^. Given that other candidate genes identified by single-cell screening, such as CYP19A1, FSHR and IL18, have already been reported in ovarian aging^[43,44]^, virtual knockout was focused on MDK and SERPINE1, whose roles remain unclear, to observe changes in the structure and strength of the regulatory network after knockout. The results showed that perturbation of MDK significantly altered genes including HP, MSLN and SERPINE2, whereas perturbation of SERPINE1 markedly changed genes including MAP3K5 and DOCK5 (Fig. 3A). These results suggest that MDK and SERPINE1 may participate in extracellular matrix homeostasis abnormality and cellular stress response, respectively, and are important candidate targets linking ovarian aging with matrix microenvironment abnormalities.

**Fig. 3.**
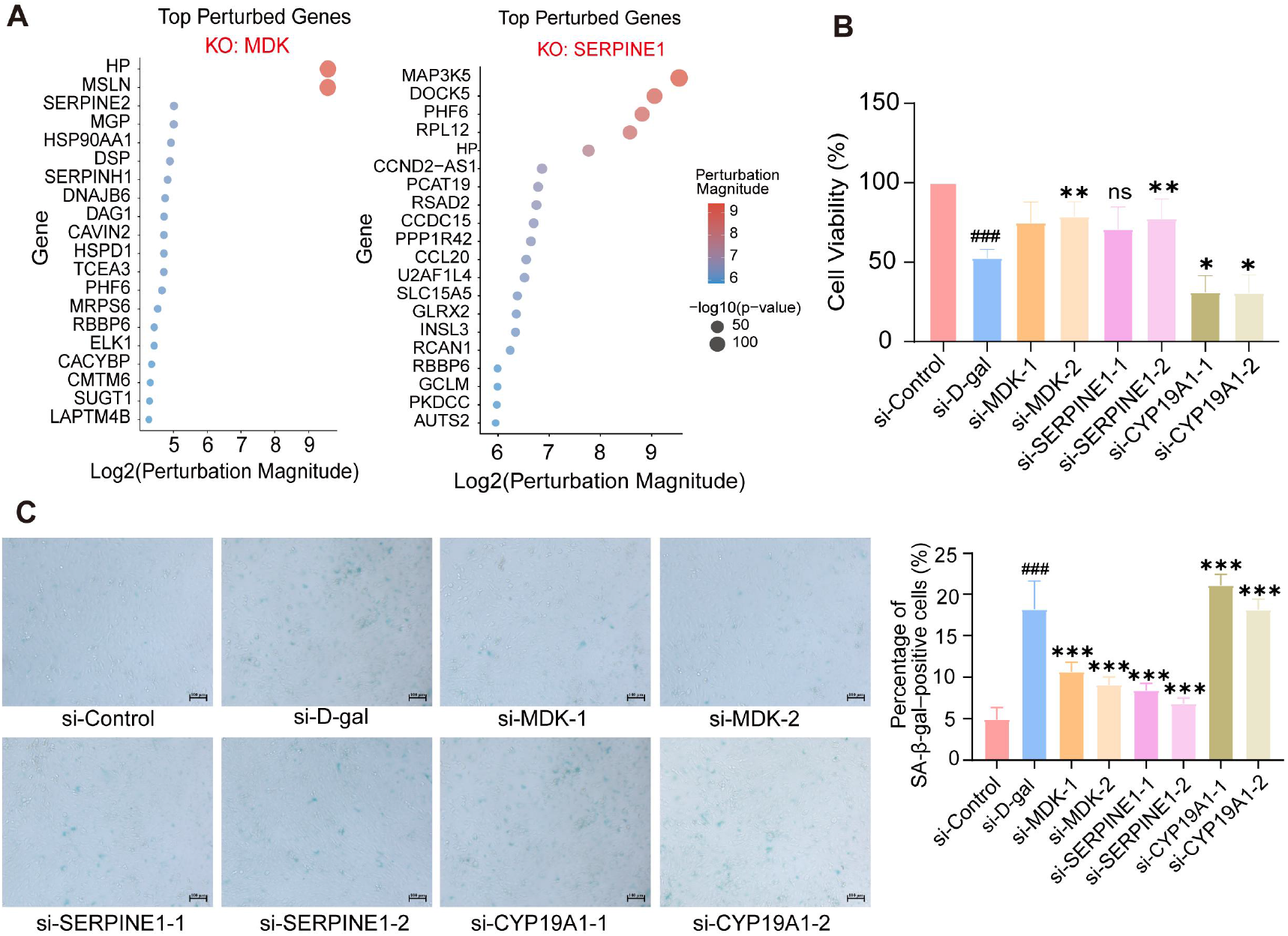
Virtual knockout and transient knockdown validate the roles of MDK, SERPINE1 and CYP19A1 in granulosa cell senescence. (A) Top perturbed genes after virtual knockout of MDK (left) and SERPINE1 (right) in granulosa cells. (B) CCK-8 cell viability in each group. (C) Representative SA-β-gal staining images and quantification.

To validate the functions of the above candidate targets experimentally, two independent siRNAs were used to transiently knock down MDK, SERPINE1 and CYP19A1, respectively; qRT-PCR and Western blot confirmed that all three genes were effectively knocked down at both the mRNA and protein levels (Fig. S3). Phenotypic assays showed that, compared with the si-D-gal group, knockdown of MDK or SERPINE1 improved the D-gal-induced decrease in cell viability and reduced the proportion of SA-β-gal-positive cells, whereas knockdown of CYP19A1 further decreased cell viability and increased the proportion of SA-β-gal-positive cells (Fig. 3B, C). These results indicate that MDK and SERPINE1 may act as pro-aging factors in ovarian aging, whereas CYP19A1 may exert anti-aging effects.

### 3.3 Human-use evidence mining identifies 10 key MFH herbs and constructs the MFH molecular library

A total of 505 prescriptions for ameliorating ovarian aging and related diminished ovarian function phenotypes were obtained, involving 295 herbs with a cumulative usage frequency of 5,803 (Fig. 4A). Among them, 69 MFH herbs accounted for 44.12% of total usage, indicating that MFH herbs have a high application weight in the relevant prescriptions. Analysis of properties, flavors and meridian tropism showed that MFH herbs were dominated by tonifying and replenishing herbs, with the four properties mainly warm and neutral, the five flavors mainly sweet, and meridian tropism centered on the kidney, liver and spleen (Fig. 4B-D). Angelicae Sinensis Radix and Rehmanniae Radix Praeparata co-occurred frequently (Fig. 4E). Association rule analysis yielded 40 valid rules, in which Rehmanniae Radix Praeparata occupied a core position in multiple high-confidence rules and formed stable combinations with Angelicae Sinensis Radix, Dioscoreae Rhizoma and Corni Fructus (Fig. 4F-G). Based on frequency, compatibility association and clustering results, 10 key MFH herbs were selected: Rehmanniae Radix Praeparata, Angelicae Sinensis Radix, Glycyrrhizae Radix et Rhizoma, Poria, Dioscoreae Rhizoma, Corni Fructus, Lycii Fructus, Ginseng Radix et Rhizoma, Astragali Radix and Rehmanniae Radix.

**Fig. 4.**
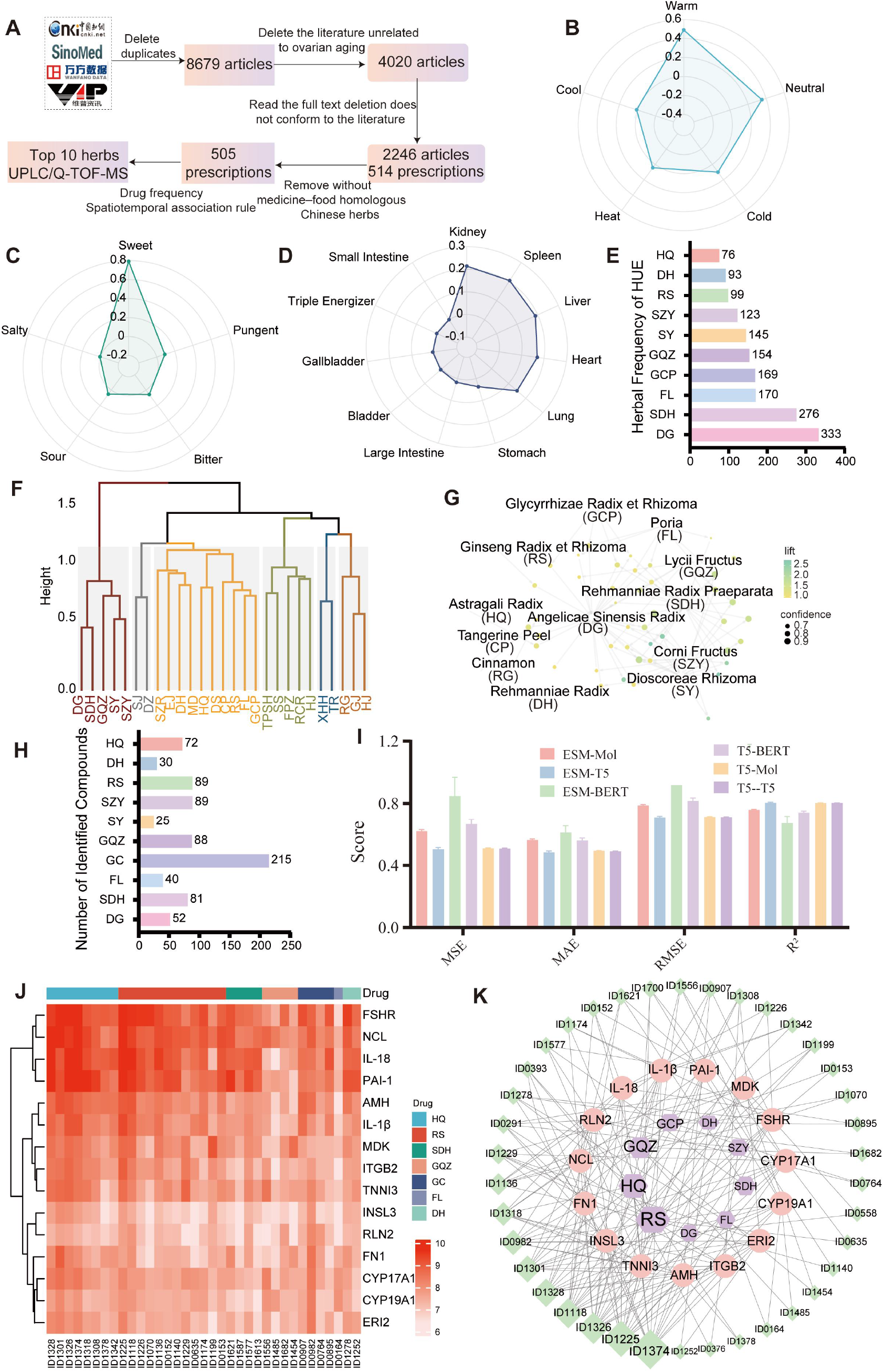
Human-use evidence mining and MFH molecular library construction for herbal candidate screening. (A) Overview of prescription mining for ovarian aging-related conditions, including the number of prescriptions, herbs and total usage frequency.(B) Distribution of herbal categories among medicine–food homology herbs.(C) Distribution of four properties and five flavors of medicine–food homology herbs. (D) Meridian tropism distribution of medicine–food homology herbs.(E) High-frequency medicine–food homology herbs with usage frequency ≥20.(F) Association rule analysis of high-frequency medicine–food homology herbs based on the Apriori algorithm. (G) Cluster analysis of core medicine–food homology herbs. (H) Construction of the MFH molecular library based on UPLC/Q-TOF-MS annotation and structural standardization. (I) ESM-T5 training results. (J) Heatmap of predicted pKD values between candidate molecules and key ovarian aging-associated targets. (K) Herb–compound–target network constructed using the top 10 predicted molecules for each key target.

Standard decoctions of the 10 key herbs were analyzed by UPLC/Q-TOF-MS, and a total of 781 compounds were identified across the herbal materials (Fig. 4H, Fig. S4); after deduplication and structural standardization, 712 small molecules suitable for subsequent prediction were obtained, constituting the MFH molecular library (Tab. S1).

### 3.4 Model combination predicts compound–target affinity and generates the core herb pair

Based on the performance comparison of six protein–molecular pretrained representation combinations under a unified evaluation framework, ESM-T5 was identified as the core configuration for subsequent compound–target affinity prediction (Fig. 4I). Replacing empirical specification with performance comparison reduces the subjectivity of model selection and improves the reproducibility of the screening workflow. Using the 712 small molecules in the MFH molecular library and 29 ovarian aging-related targets identified by single-cell analysis, ESM-T5 was applied to predict compound–target affinity, constructing a 712 × 29 prediction matrix. The top 10 molecules ranked by predicted pK_D for each target were extracted for visualization, and multiple small molecules derived from Astragalus and ginseng showed relatively high predicted affinity for several key targets (Tab. S2).

A herb–compound–target network was constructed based on the top 10 molecules ranked by predicted pK_D for each target. The numbers of high-affinity molecules derived from Astragalus and ginseng were relatively large, at 9 and 12 respectively, and their target coverage ranked highest (Fig. 4J, K). Considering the enrichment of high-affinity molecules, target coverage and network topological features, the Astragalus–ginseng combination (HXF) was finally identified as the core candidate herb pair for subsequent in vitro and in vivo validation.

### 3.5 Molecular dynamics simulation and SPR assays evaluate complex stability

To verify the reliability of affinity prediction at the computational level, high-affinity candidate compounds were selected for representative targets including CYP19A1, PAI-1, MDK, NCL, IL18, ITGB2 and AMH to construct complexes, and 100 ns molecular dynamics (MD) simulations were performed. The results showed that the binding free energies of most systems were below −7 kcal/mol. The RMSD of each complex increased during 0–20 ns to accommodate ligand binding and mostly stabilized after approximately 40 ns; only ITGB2 fluctuated markedly throughout, indicating weaker binding stability. Except for MDK, which showed high peaks in some regions and relatively large local flexibility, most residues of the other complexes had RMSF < 0.2 nm. All systems maintained a certain number of hydrogen bonds throughout the simulation, with the AMH-, CYP19A1– and PAI-1-related complexes showing more numerous and stable hydrogen bonds (Fig. 5A–C, Fig. S5). Conformational analysis further showed that most complexes maintained stable binding conformations within 100 ns, with ligands remaining in the binding pocket throughout (Fig. S6).

**Fig. 5.**
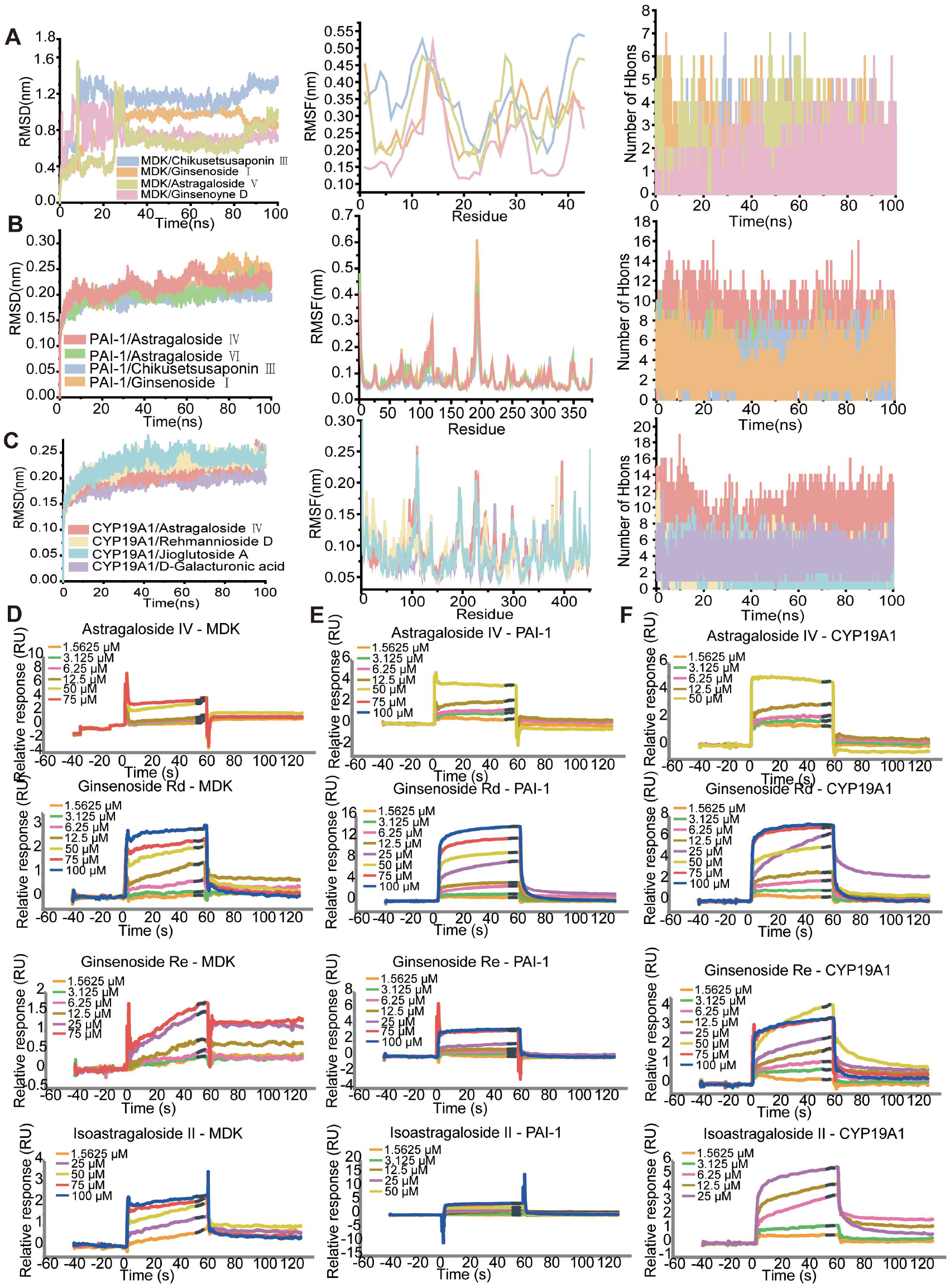
Molecular dynamics simulation and surface plasmon resonance (SPR) validation of key molecule–target complexes. (A–C) The 100 ns molecular dynamics simulation results for MDK, PAI-1, and CYP19A1 in complex with representative active components, with the three panels in each row showing, from left to right, the root mean square deviation (RMSD), residue root mean square fluctuation (RMSF), and the number of hydrogen bonds. (D–F) SPR binding curves and fitted K_D values for the interactions of Astragaloside IV, Ginsenoside Rd, Ginsenoside Re, and Isoastragaloside II with MDK, PAI-1, and CYP19A1, respectively.

Based on the ESM-T5 affinity prediction results, top-ranked compounds, namely ginsenoside Rd, ginsenoside Re, isoastragaloside II and astragaloside IV, were selected and paired with CYP19A1, PAI-1 and MDK for SPR validation. Of the 12 pairs, 10 were successfully fitted, and molecule–target combinations with fittable parameters yielding K_D_ values accounted for 83.33%. Fittable results were obtained for ginsenoside Rd, ginsenoside Re, isoastragaloside II and astragaloside IV, with K_D_ values of 58.9, 57.3, 58.0 and 22.1 µM, respectively, indicating good consistency between ESM-T5 predictions and experimental measurements (Fig. 5D–E). SPR and MD simulations corroborate each other, verifying from both computational and experimental perspectives that some high-affinity molecule–target combinations predicted by ESM-T5 possess detectable binding activity.

### 3.6 Quality control of HXF and its amelioration of the D-gal-induced senescence phenotype and related molecular expression in KGN granulosa cells

To ensure the reproducibility of subsequent pharmacological experiments, representative ginseng saponins (ginsenoside Re and Rd) and characteristic Astragalus constituents (astragaloside IV and isoastragaloside II) were used as quality control markers for HXF, and their contents were determined by HPLC-ELSD. Standard curves were plotted using the four reference standards, and method validation was performed for linearity, precision, LOD and LOQ (Fig. S7). The contents of the four marker constituents were similar across three batches of HXF test samples, indicating good batch-to-batch consistency (Fig. 6A).

**Fig. 6.**
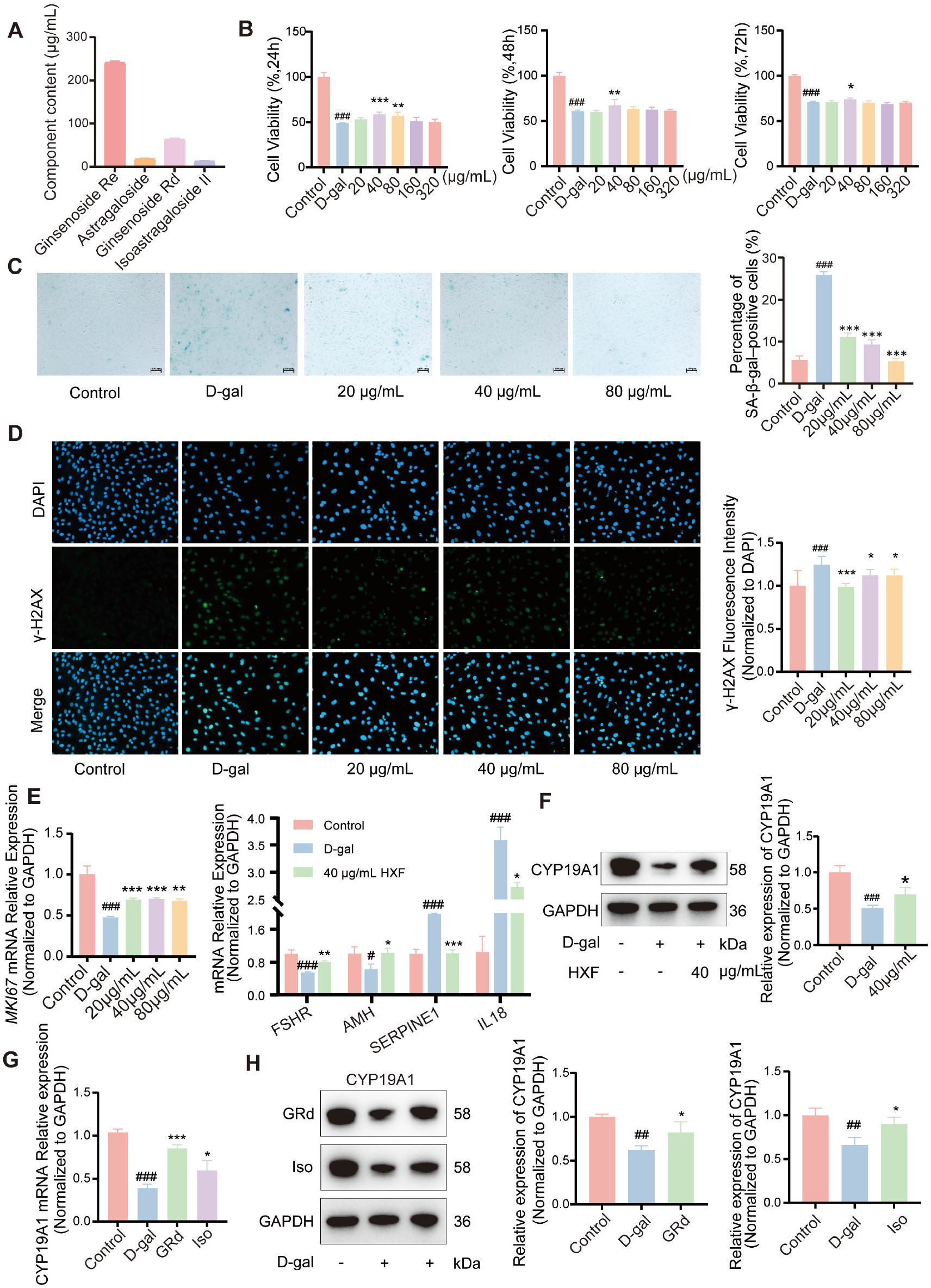
HXF attenuates D-gal-induced granulosa cell senescence and restores functional marker expression. (A) Contents of representative marker compounds in HXF. (B) Effects of HXF on the viability of D-gal-treated KGN cells assessed by CCK-8 assay. (C) Representative images and quantitative analysis of SA-β-gal staining in KGN cells. (D) Representative immunofluorescence images and quantification of γ-H2AX in KGN cells.(E) qPCR and Western blot analysis of FSHR, AMH, MKI67, SERPINE1 and IL18 after D-gal and HXF treatment. (F) Western blot analysis of CYP19A1 protein expression in KGN cells. (G) qPCR analysis of CYP19A1 mRNA expression after D-gal, Ginsenoside Rd (GRd), and Isoastragaloside II (Iso) treatment. (H) Western blot analysis of CYP19A1 protein expression in KGN cells after D-gal, Ginsenoside Rd (GRd), and Isoastragaloside II (Iso) treatment.

CCK-8 assays showed that D-gal significantly reduced cell viability (P < 0.001), whereas HXF intervention restored viability to variable degrees. The effects of 40 and 80 µg/mL were statistically significant at 24, 48 and 72 h, with 40 μg/mL for 24 h yielding the best effect (P < 0.001); this condition was therefore adopted for subsequent experiments (Fig. 6B). Phenotypic assays showed that, relative to the D-gal group, HXF intervention reduced the SA-β-gal-positive rate and γ-H2AX level (Fig. 6C–D). HXF upregulated the mRNA expression of the KGN functional molecules MKI67, FSHR and AMH and the protein expression of CYP19A1, while downregulating the mRNA levels of the senescence-associated molecules SERPINE1 and IL18 (Fig. 6E–F). Ginsenoside Rd and Isoastragaloside II increased CYP19A1 expression at both the mRNA and protein levels (Fig. 6G–H). Taken together, HXF alleviates the D-gal-induced cellular senescence phenotype by preserving granulosa cell function, promoting proliferation and suppressing senescence-related inflammation, and its regulatory trend is consistent with the key targets predicted by single-cell transcriptomic analysis.

### 3.7 HXF ameliorates ovarian function impairment and endocrine disorder in CTX-induced ovarian aging rats

Rats in the model group showed reduced body weight and decreased ovarian index, together with increased liver, spleen and kidney indices, which were reversed to varying degrees after 2 weeks of HXF intervention (Fig. 7A, B; Fig. 8A). Estrous cycle and ovarian histomorphological examinations showed that HXF at all doses reduced the proportion of CTX-induced estrous cycle abnormalities (Fig. 8B) and significantly decreased the proportion of atretic follicles (P < 0.05), indicating that HXF alleviated ovarian tissue structural damage and improved follicular development (Fig. 7C, D). Immunohistochemistry showed that, compared with the model group, HXF upregulated KI67 expression (8.4 g/kg, P < 0.01) and reduced the γ-H2AX-positive area (4.2 and 8.4 g/kg, P < 0.05) (Fig. 8E, F). Serum measurements showed that E2 was significantly decreased in the model group, whereas the FSH level and the FSH/LH ratio were significantly increased; HXF at all doses restored E2 (P < 0.05), and 4.2 and 8.4 g/kg also reduced FSH and the FSH/LH ratio (P < 0.05), whereas AMH showed no significant difference among groups (Fig. 7G). Therefore, HXF ameliorated body weight loss and ovarian tissue structure, restored the estrous cycle, promoted ovarian cell proliferation and alleviated DNA damage, while reducing serum sex hormone disturbance in CTX-induced ovarian aging rats.

**Fig. 7.**
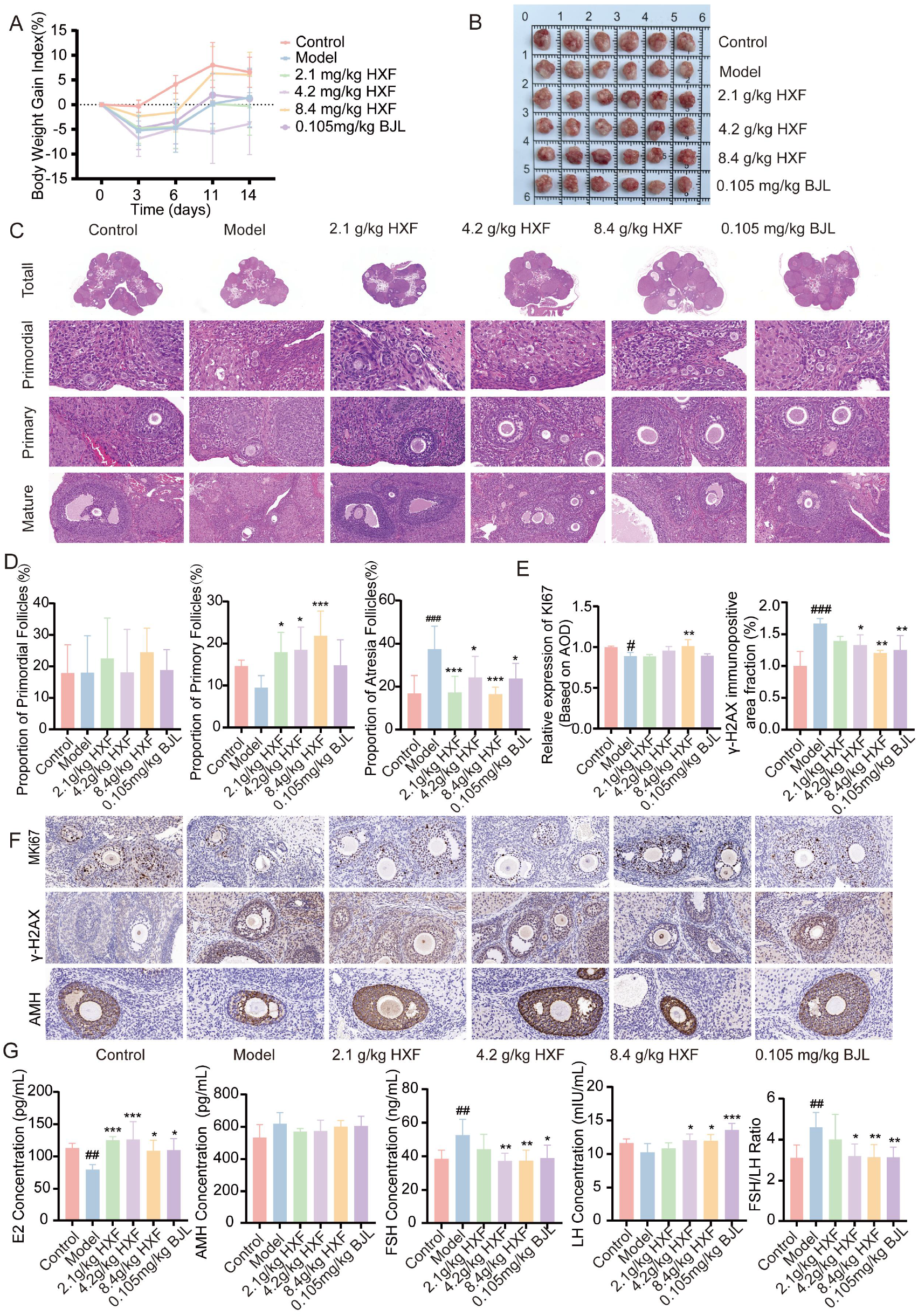
HXF ameliorates CTX-induced ovarian injury and sex hormone secretion in Ovarian ageing rats. (A) Changes in the body weight gain index of rats in each group. (B) Representative gross morphology of ovaries from each group. (C) Representative H&E-stained ovarian sections showing the overall ovarian architecture (Total) and follicles at different developmental stages (primordial, primary, and mature follicles); scale bar = 100 μm. (D) Quantitative analysis of the proportions of primordial, primary, and atretic follicles. (E) Quantitative analysis of KI67 expression and the γ-H2AX-immunopositive area fraction. (F) Representative immunohistochemical images of KI67, γ-H2AX, and AMH in ovarian tissues. (G) Serum levels of E2, AMH, FSH, and LH, and the FSH/LH ratio in each group.

**Fig. 8.**
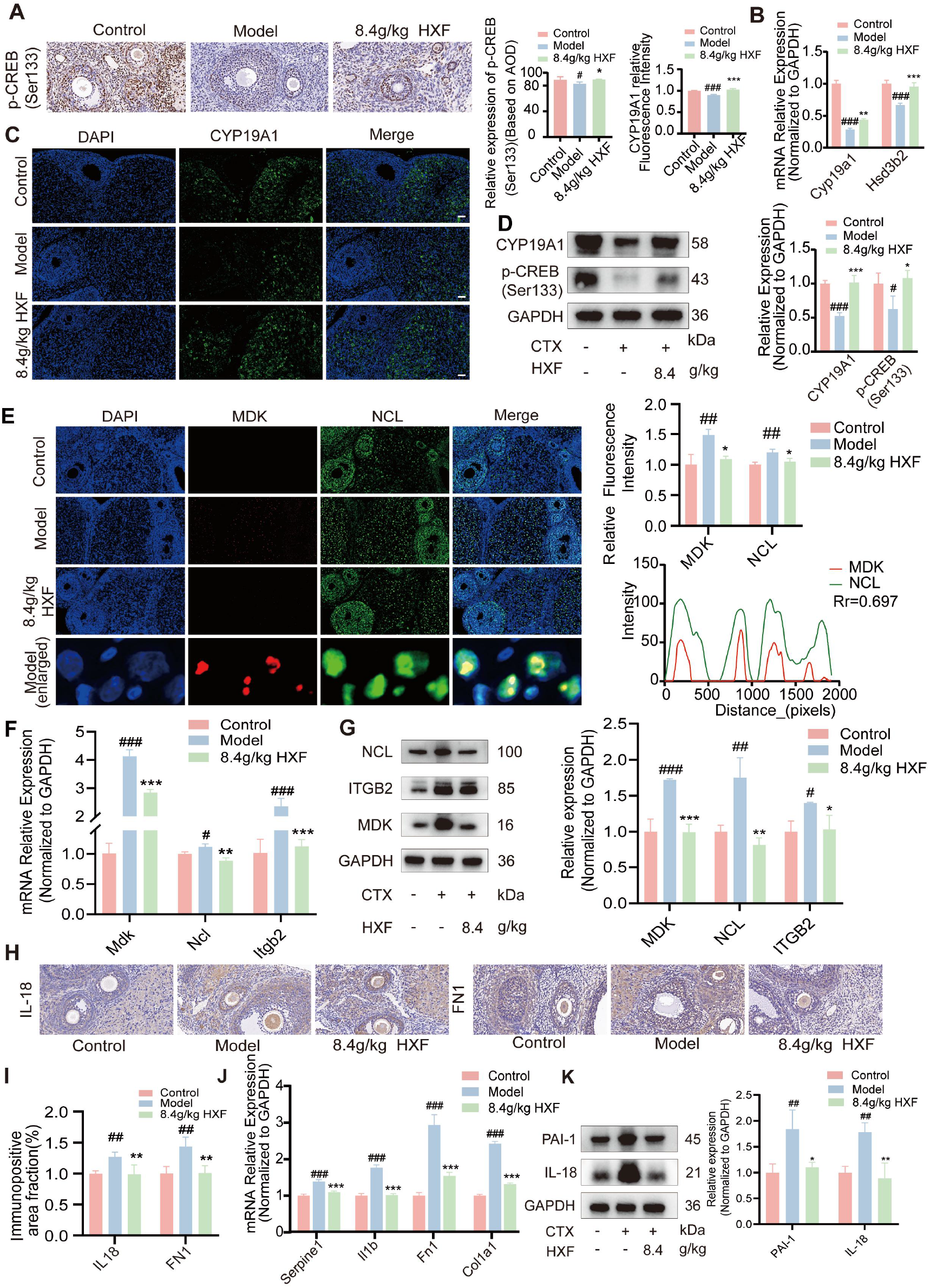
HXF ameliorates CTX-induced steroidogenic dysfunction in Ovarian ageing rats and modulates MDK–NCL signaling as well as inflammation– and extracellular matrix remodeling-related molecules. (A) Representative immunohistochemical staining images of p-CREB in ovarian tissue (200×) and quantitative analysis; (B) relative mRNA expression levels of Cyp19a1 and Hsd3b2 in ovarian tissue; (C) representative immunofluorescence images of CYP19A1 (200×) and quantitative analysis; (D) Western blot bands and densitometric quantification of CYP19A1 and p-CREB (Ser133); (E) double-label immunofluorescence staining images of MDK and NCL; (F) relative mRNA expression levels of Mdk, Ncl and Itgb2; (G) Western blot bands and densitometric quantification of NCL, ITGB2 and MDK; (H) representative immunohistochemical staining images of IL-18 and FN1 (200×); (I) quantification of the immunopositive area fraction (%) of IL-18 and FN1; (J) relative mRNA expression levels of Serpine1, Il1b, Fn1 and Col1a1; (K) Western blot bands and densitometric quantification of PAI-1 and IL-18.

### 3.8 HXF ameliorates steroidogenic dysfunction and modulates MDK–NCL communication and inflammation– and extracellular matrix remodeling-related molecules in CTX-induced ovarian aging rats

Transcriptome sequencing of ovarian tissue showed that PCA clearly separated the three groups of samples in principal component space, with the 8.4 g/kg HXF group shifted relative to the model group. Venn analysis identified 101 common differentially expressed genes, of which 29 were upregulated and 34 downregulated in the model group (Fig. S9). Enrichment analysis showed that the differentially expressed genes were mainly enriched in inflammation– and stress-related pathways such as TNF, IL-17, MAPK and cytokine–receptor interaction, and also involved the Hippo and ovarian steroidogenesis pathways, with cellular components centered on the extracellular region and extracellular matrix (Fig. S10). Based on these findings, validation of the effect of HXF on the ovarian steroidogenesis pathway showed that Cyp19a1 and Hsd3b2 mRNA as well as p-CREB (Ser133) and CYP19A1 protein expression were significantly downregulated in the model group, and all these indicators were significantly increased after HXF intervention (Fig. 8A-D). Given that single-cell communication analysis indicated involvement of the MDK–NCL axis in aberrant cell communication during ovarian aging, this was validated in the CTX model. The mRNA and protein expression of Mdk, Ncl and Itgb2 were significantly upregulated in the model group and downregulated after HXF intervention; immunofluorescence showed obvious colocalization of MDK and NCL, with enhanced fluorescence signals in the model group and attenuated signals in the HXF group (Fig. 8E-G). Given that diminished ovarian function is often accompanied by inflammatory activation and extracellular matrix remodeling, this study found that Serpine1, Col1a1, Il1b and Fn1 mRNA as well as IL-18, FN1 and PAI-1 protein expression were significantly increased in the model group, and all significantly decreased toward normal levels after HXF intervention (Fig. 8H-K). Therefore, HXF significantly ameliorated CTX-induced diminished ovarian function, involving remodeling of the ovarian transcriptional regulatory network, restoration of the CYP19A1/HSD3B2-mediated steroidogenesis pathway, suppression of aberrant activation of the MDK–NCL signaling axis, and improvement of the ovarian microenvironment and promotion of tissue function recovery by alleviating inflammatory responses and correcting SERPINE1/PAI-1-related fibrinolytic system abnormalities and extracellular matrix remodeling.

## 4. Discussion

Ovarian aging is an important biological basis for female reproductive aging and increased life-course health risks. its onset and progression involve multi-level abnormalities including follicular reserve depletion, granulosa cell functional decline, steroidogenic dysfunction, enhanced inflammatory responses and ovarian microenvironment remodeling^[45,46]^. Although traditional Chinese herbal formulas have long-standing human-use experience, their screening relies largely on empirical summarization, frequency statistics and network association analysis, making it difficult to incorporate disease mechanisms and compound–target relationships^[47]^. Advances in databases and AI algorithms have improved research efficiency, but have still not focused on the integration of disease mechanisms, compound–target relationships and experimental validation in formula screening^[48]^. Taking ovarian aging as the scenario, this study constructed an MFH herbal combination discovery workflow running through disease mechanism decoding, human-use evidence constraint, pretrained representation combination screening and multi-level experimental validation. Rather than redeveloping large-scale pretrained models, this workflow reuses existing protein and molecular representations through modular integration and unified evaluation, selects the affinity prediction configuration suitable for the present task, and completes closed-loop evaluation of candidate herb pairs by combining human-use evidence with experimental validation, ultimately identifying Astragalus–ginseng as the candidate herb pair.

Based on single-cell transcriptomic analysis, this study showed that aged ovaries exhibit not only abnormalities in steroidogenesis– and follicular function-related molecules such as CYP19A1, HSD3B2 and FSHR, but also enhanced immune cell communication, activated inflammatory signaling and extracellular matrix remodeling, consistent with the view that ovarian aging involves global microenvironmental imbalance rather than a single endocrine decline^[49,50]^. Among the candidate targets, SERPINE1 was markedly upregulated in granulosa cells and vascular endothelial cells; its encoded protein PAI-1 participates not only in fibrinolytic regulation but is also closely related to cellular senescence, inflammatory amplification and tissue fibrosis^[51–54]^, indicating that fibrinolytic homeostasis imbalance and matrix remodeling are important components of ovarian aging. The altered expression of stress– and migration-related genes such as MAP3K5 and DOCK5 revealed by virtual knockout supports its involvement in granulosa cell stress injury and microenvironmental abnormalities^[55]^. Cell communication analysis revealed enhanced interaction of the MDK–NCL ligand–receptor axis between granulosa cells and multiple cell types. MDK is a secreted factor involved in development, tissue repair, inflammation and age-related diseases, whereas NCL can serve as its cell-surface binding molecule mediating signal transduction; the MDK–NCL axis can drive immunosuppression and induce M2 polarization of macrophages in various pathological processes^[56,57]^. In this study, the downstream gene profiles after perturbation of the two were highly similar, involving adhesion-, fibrinolysis– and matrix homeostasis-related molecules such as MSLN, SERPINE2 and SERPINH1^[58–60]^. Moreover, MDK can enhance neutrophil adhesion by promoting the high-affinity conformation of β2 integrin (ITGB2), supporting immune cell trafficking during acute inflammation^[61]^. Combined with the hub positions of FN1 and IL1B in the PPI network and the changes in ITGB2 and IL18, this study proposes that ovarian aging involves an interconnected pathological network of steroidogenic dysfunction, aberrant cell communication, inflammatory amplification and fibrinolysis/matrix remodeling.

At the level of pretrained model screening, this study highlights the adaptation problem between the rapid iteration of pretrained models and their implementation in TCM research and development. Retraining large-scale models imposes high requirements on data, computing power and algorithmic expertise^[17]^. Existing platforms mostly adopt fixed or closed architectures, making it difficult to integrate new model resources in a timely manner^[19]^. To address this, this study treated protein and molecular representation models as replaceable modules and compared them in combination under a unified dataset, fusion strategy, downstream regression architecture and evaluation metrics. ESM-T5 showed relatively better overall performance under the data and task conditions of this study and was subsequently used for compound–target affinity prediction on the MFH molecular library. Notably, the performance differences among ESM-T5 and combinations such as T5–T5 and T5–Mol were not substantial. This indicates that under the current data scale, feature concatenation strategy and downstream regression architecture, the advantages of different pretrained representation combinations are not absolute^[62]^. It should therefore be understood as a relative preference under the conditions of this task rather than a universal optimum; the value of this strategy lies in providing a low-cost workflow for reusing, evaluating and updating existing pretrained representation resources, rather than proposing a new model.

In terms of herbal combination screening strategy, this study did not adopt a fully open-ended natural product virtual screening approach; instead, human-use evidence for medicine–food homology herbs was used as a prior constraint to construct the candidate herbal material and molecular libraries. MFH herbs possess both edible and medicinal attributes with a favorable safety foundation, making them suitable for long-term intervention in chronic, progressive ovarian aging^[63,64]^. Mining of literature and classical prescriptions revealed that MFH herbs have a high application weight in prescriptions related to ovarian aging, with medicinal properties concentrated in tonifying deficiency, sweet flavor, warm and neutral nature, and tropism toward the kidney, liver and spleen meridians, consistent with the TCM understanding that kidney deficiency is the root accompanied by qi and blood disharmony^[65–67]^. On this basis, UPLC/Q-TOF-MS chemical profiling, ESM-T5 affinity prediction, molecular dynamics simulation, SPR and network analysis were integrated to identify Astragalus–ginseng as the core herb pair. This result was jointly supported by human-use evidence, compound detection, target coverage, enrichment of high-affinity molecules and experimental validation, rather than determined solely by frequency or computed affinity, thereby improving the accuracy of the screening results.

Both Astragalus and ginseng are representative qi-tonifying herbs containing multiple classes of constituents such as saponins, polysaccharides and flavonoids, with immunomodulatory, antioxidant and tissue-protective effects^[68]^. Modern pharmacological studies also indicate that both contain multiple classes of active constituents including saponins, polysaccharides and flavonoids, with effects of regulating immunity, antioxidation, improving energy metabolism and tissue protection^[69,70]^. In this study, astragaloside IV, isoastragaloside II, ginsenoside Rd and ginsenoside Re showed relatively high affinity for PAI-1, CYP19A1 or MDK, providing a preliminary material basis for the action of HXF on key constituents. However, SPR covered only a subset of representative molecules and targets; the effects of herbal combinations are more likely to arise from the synergistic regulation of multiple constituents across multiple pathway nodes^[71]^, and the key pharmacodynamic substances remain to be clarified by serum pharmacochemistry, pharmacokinetics and component deconvolution experiments.

Granulosa cells are the principal cells involved in ovarian steroidogenesis and also serve as sensors and regulators of microenvironmental changes; their functional state directly determines follicular fate and ovarian reserve^[72,73]^. In vitro experiments showed that HXF increased the viability of D-gal-induced senescent KGN cells, reduced the SA-β-gal-positive rate and γ-H2AX signal, restored the expression of FSHR, AMH, MKI67 and CYP19A1, and downregulated SERPINE1 and IL18, indicating that it can act directly on granulosa cells to alleviate senescence, DNA damage and functional decline.

Recent studies have found that ovarian aging manifests not only as a reduction in follicle number, but also as enhanced chronic low-grade inflammation, altered immune cell composition, aggravated tissue fibrosis and local microenvironmental imbalance^[74,75]^. Although the CTX-induced ovarian aging rat model is not equivalent to natural age-related ovarian aging, the two share high similarity in several key phenotypes^[76,77]^. This study showed that HXF improved estrous cycle disturbance, decreased ovarian index, impaired follicular development and abnormal serum FSH and E2, reduced γ-H2AX and increased KI67, while upregulating Cyp19a1, Hsd3b2, p-CREB (Ser133) and CYP19A1, suggesting that it may improve ovarian endocrine function by restoring the CREB-related steroidogenesis pathway. In addition, HXF downregulated the expression of Mdk, Ncl and Itgb2, attenuated the colocalization signal of MDK and NCL in ovarian tissue, and reduced the levels of inflammation-, fibrinolytic system– and extracellular matrix remodeling-related molecules such as Serpine1, Il1b, Il18, Fn1, Col1a1 and PAI-1. These results corroborate the single-cell analysis and constitute an evidence loop from computational screening to experimental validation. Causal validation is ongoing.

This study still has several limitations. First, the evaluation of pretrained representation combinations was mainly based on ChEMBL and BindingDB data; although five-fold cross-validation was performed, external independent validation and cold-start scenario evaluation are still lacking, and future work could examine model generalization to new scaffold compounds and new targets. Second, the current model fusion is mainly based on feature concatenation, which insufficiently models fine-grained cross-modal interactions between proteins and ligands; attention mechanisms, graph neural networks or multi-model ensemble methods could be introduced to improve prediction performance. Third, the material basis of the Astragalus–ginseng herb pair has not been fully elucidated, and SPR and MD experiments covered only a subset of representative constituents; in vivo experiments are still needed to confirm the key pharmacodynamic constituents. Fourth, the CTX model cannot fully simulate the natural ovarian aging process; future studies should combine natural aging models and primary cell systems for multi-model validation. Fifth, validation of MDK–NCL, SERPINE1/PAI-1 and CYP19A1 mainly remained at the level of molecular expression, and functional intervention experiments are still needed to clarify causal roles.

In summary, this study constructed an MFH herbal combination screening workflow that integrates single-cell mechanism decoding, human-use evidence constraints, modular pretrained representation combinations and multi-level experimental validation, and completed a closed-loop application in the ovarian aging scenario, from key target identification and candidate herb pair screening and quality control to cellular and animal experimental validation. The results show that the Astragalus–ginseng combination can ameliorate ovarian aging-related phenotypes to a certain extent, and its effects may be related to restoring CYP19A1-mediated steroidogenesis, modulating aberrant MDK–NCL cell communication, and alleviating SERPINE1/IL18-related inflammatory responses and extracellular matrix remodeling. This provides a methodological reference for the mechanism-oriented screening of MFH herbal combinations and for the low-cost reuse of pretrained model resources in TCM research and development.

## 5. Conclusion

This study established an integrated workflow combining single-cell mechanism decoding, human-use evidence constraints, UPLC/Q-TOF-MS chemical profiling, evaluation of pretrained protein and molecular representation combinations, compound–target affinity prediction, and in vitro and in vivo experimental validation. In the ovarian aging scenario, the Astragalus–ginseng core herb pair was identified, and in vitro and in vivo experiments confirmed that it ameliorated KGN cell senescence and CTX-induced ovarian aging in rats. Its effects involve restoring CYP19A1/HSD3B2-mediated steroidogenesis, suppressing aberrant MDK–NCL axis cell communication, and alleviating SERPINE1/IL18-related inflammatory responses and extracellular matrix remodeling. By replacing repeated model development with modular integration, this workflow provides a reusable and updatable methodological reference for the mechanism-oriented screening of MFH herbal combinations.

### CRediT authorship contribution statement

Jinmei Zhang: Writing manuscript, data curation and analysis, investigation. Xia Chen: Single-cell sequencing data analysis, in vitro and in vivo experiments. Yahui Cao: Model construction and prediction. Yaou Liu: Animal histopathology experiments. Nan Ning: Human-use evidence collection. Weijie Bian: SPR experiments. Xiangning Yang: Molecular dynamics simulation. Yanlei Guo: Chemical profiling. Jin Zeng: Overall experimental design, project administration, manuscript review.

### Ethical approval

The SD rats used in the animal experiments were purchased from Beijing Huafukang Bioscience Co., Ltd., laboratory animal production license No. SCXK (Jing) 2024-0003. They were housed in the SPF animal experiment center of the Sichuan Academy of Chinese Medicine Sciences, laboratory animal use license No. SYXK (Chuan) 2023-0100. All animal experimental procedures followed the regulations of the Animal Welfare and Ethics Committee of the Sichuan Academy of Chinese Medicine Sciences and were approved by that committee, with ethics approval No. DWSYLL-2025-100.

### Data availability statement

The data, materials and experimental protocols involved in this study are available from the corresponding author upon reasonable request, subject to journal policies and relevant ethical and legal requirements.

### Funding

This work was supported by the Science and Technology Research Special Project of the Sichuan Provincial Administration of Traditional Chinese Medicine, China (25ZDIZX013), the General Program of the National Natural Science Foundation of China (82675540), and the Project of the Science and Technology Department of Sichuan Province, China (2025JDKY0027).

## Acknowledgements

The authors thank the Sichuan Academy of Chinese Medicine Sciences, Tianjin University and the Peking University Chengdu Academy for Advanced Interdisciplinary Biotechnologies for providing instrument platforms and technical support.

## Supplementary material

Supplementary material associated with this article can be found in the online version.

## References

[1] Ruth K S, Day F R, Hussain J, et al. Genetic insights into biological mechanisms governing human ovarian ageing[J]. Nature, 2021, 596(7872): 393–397. DOI: 10.1038/s41586-021-03779-7.

[2] Secomandi L, Borghesan M, Velarde M, et al. The role of cellular senescence in female reproductive aging and the potential for senotherapeutic interventions. Hum Reprod Update. 2022, 28(2):172–189. DOI: 10.1093/humupd/dmab038.

[3] Dong L, Teh D B L, Kennedy B K, et al. Unraveling female reproductive senescence to enhance healthy longevity[J]. Cell Research, 2023, 33(1): 11–29. DOI: 10.1038/s41422-022-00718-7.

[4] Kerr J B, Rodgers R J. The human ovarian reserve: The narrative and the science[J/OL]. Human Reproduction Update, 2025: dmaf031. DOI: 10.1093/humupd/dmaf031.

[5] Liang X, Yan Z, Ma W, et al. Peroxiredoxin 4 protects against ovarian ageing by ameliorating d-galactose-induced oxidative damage in mice[J]. Cell Death & Disease, 2020, 11(12): 1053. DOI: 10.1038/s41419-020-03253-8.

[6] Mehta J M, Manson J E. The menopausal transition period and cardiovascular risk[J]. Nature Reviews Cardiology, 2024, 21(3): 203–211. DOI: 10.1038/s41569-023-00926-7.

[7] Branyan T E, Aleksa J, Lepe E, et al. The aging ovary impairs acute stroke outcomes[J]. Journal of Neuroinflammation, 2023, 20(1): 159. DOI: 10.1186/s12974-023-02839-1.

[8] Stankovic S, Shekari S, Huang Q Q, et al. Genetic links between ovarian ageing, cancer risk and de novo mutation rates[J]. Nature, 2024, 633(8030): 608–614. DOI: 10.1038/s41586-024-07931-x.

[9] China Association of Chinese Medicine. Integrated Chinese and Western Medicine Guidelines for Premature Ovarian Failure[J]. Chinese Journal of Integrated Traditional and Western Medicine, 2024, 44(10): 1167–1179.

[10] Cheng J M, Xue F, Zhang Y Y, et al. Research and Development Status, Technical Keys and Countermeasures of Medicinal and Food Homologous Products[J]. Journal of Nanjing University of Traditional Chinese Medicine, 2023, 39(9): 814–826. DOI: 10.14148/j.issn.1672-0482.2023.0814.

[11] Papalexi E, Satija R. Single-cell RNA sequencing to explore immune cell heterogeneity[J]. Nature Reviews Immunology, 2018, 18(1): 35–45. DOI: 10.1038/nri.2017.76.

[12] Elnaggar A, Heinzinger M, Dallago C, et al. ProtTrans: Toward Understanding the Language of Life Through Self-Supervised Learning[J]. in IEEE Transactions on Pattern Analysis and Machine Intelligence, 2022, 44(10): 7112–7127. DOI: 10.1109/TPAMI.2021.3095381.

[13] Rives A, Meier J, Sercu T, et al. Biological structure and function emerge from scaling unsupervised learning to 250 million protein sequences[J]. Proceedings of the National Academy of Sciences of the United States of America, 2021, 118(15). DOI: 10.1073/pnas.2016239118.

[14] Zheng X, Tomiura Y. A BERT-based pretraining model for extracting molecular structural information from a SMILES sequence[J]. Journal of Cheminformatics, 2024, 16(1): 71. DOI: 10.1186/s13321-024-00848-7.

[15] Edwards C, Lai T, Ros K, et al. Translation between molecules and natural language[J]. Proceedings of the 2021 Conference on Empirical Methods in Natural Language Processing, 2022. DOI: 10.18653/v1/2022.emnlp-main.26.

[16] Ross J, Belgodere B, Chenthamarakshan V, et al. Large-scale chemical language representations capture molecular structure and properties[J]. Nature Machine Intelligence, 2022, 4(12): 1256–1264. DOI: 10.1038/s42256-022-00580-7.

[17] Han F, Li Y, Chen Z, et al. Key technologies of AI large language models in the field of traditional chinese medicine international journal of pattern recognition and artificial intelligence[J]. International Journal of Pattern Recognition and Artificial Intelligence, 2026, (prepublish): DOI:10.1142/S0218001426590263.

[18] Jia Y, Ji X, Wang X, et al. Qibo: a large language model for traditional chinese medicine[J]. Expert Systems with Applications, 2025, 284: 127672. DOI: 10.1016/j.eswa.2025.127672.

[19] Perricone U. Development and optimisation of computational tools for drug discovery[D]. Nova Science Publishers (Nova Science Publishers, Inc.), http://hdl.handle.net/10447/220600.

[20] Jin C, Wang X, Yang J, et al. Molecular and genetic insights into human ovarian aging from single-nuclei multi-omics analyses[J]. Nature Aging, 2025, 5(2): 275–290. DOI: 10.1038/s43587-024-00762-5.

[21] Wu M, Tang W, Chen Y, et al. Spatiotemporal transcriptomic changes of human ovarian aging and the regulatory role of FOXP1[J]. Nature Aging, 2024, 4(4): 527–545. DOI: 10.1038/s43587-024-00607-1.

[22] Magalhães J P. HAGR: the Human Ageing Genomic Resources[J]. Nucleic Acids Research, 2004, 33(Database issue): D537–D543. DOI:10.1093/nar/gki017.

[23] Tăcutu R, Thornton D, Johnson E, et al. Human Ageing Genomic Resources: new and updated databases[J]. Nucleic Acids Research, 2017, 46(1): 1083–1090. DOI:10.1093/nar/gkx1042.

[24] Osorio D, Zhong Y, Li G, et al. scTenifoldNet: a machine learning workflow for constructing and comparing transcriptome-wide gene regulatory networks from single-cell data[J]. Patterns, 2020, 1(9): 100139. DOI: 10.1016/j.patter.2020.100139.

[25] Zhang S, Xu Y, Zhang J, et al. Hydroxytyrosol enhances the Nrf2/HO-1 signalling pathway to inhibit oxidative stress and apoptosis and improve premature ovarian insufficiency In vitro and In vivo[J]. International journal of molecular sciences, 2026, 27(11): 4845. DOI: 10.3390/ijms27114845.

[26] Chinese Pharmacopoeia Commission. Pharmacopoeia of the People’s Republic of China, Volume I, 2025 edition [M]. Beijing: China Medical Science Press, 2025.

[27] Zhong G S. Science of Chinese Materia Medica [M]. Beijing: China Press of Traditional Chinese Medicine, 2016.

[28] Riedl M, Mukherjee S, Gauthier M. Descriptor-free deep learning QSAR model for the fraction unbound in human plasma[J]. Molecular Pharmaceutics, 2023, 20(10): 4984–4993. DOI: 10.1021/acs.molpharmaceut.3c00129.

[29] Kovacs C, Nowak L, Horvath H. Deep Learning in Structure–Activity Relationship Modeling[J]. Zenodo (CERN European Organization for Nuclear Research), 2023. DOI:10.5281/zenodo.19511647.

[30] Cho K, Edwards C, Honke G, et al. Translation between Molecules and Natural Language[J]. Underline Science Inc., 2022. DOI:10.48448/asve-g660.

[31] Liu X, Zhang R, Li J, et al. Characterization and clustering of immunocompetent proteins based on ESM-2 sequence embedding and physicochemical feature fusion[J]. Frontiers in Immunology, 2026, 17: 1918918. DOI: 10.3389/fimmu.2026.1918918.

[32] Marquet C, Heinzinger M, Olenyi T, et al. Embeddings from protein language models predict conservation and variant effects[J]. Human Genetics, 2022, 141(10): 1629–1647. DOI: 10.1007/s00439-021-02411-y.

[33] Gaulton A, Hersey A, Nowotka M, et al. The ChEMBL database in 2017[J]. Nucleic Acids Research, 2016, 45(D1): D945–D954. DOI: 10.1093/nar/gkw1074.

[34] Jin Y, Wang R, Zhuang X, et al. Prediction of COVID-19 data using an ARIMA-LSTM hybrid forecast model[J]. Mathematics, 2022, 10(21): 4001. DOI: 10.3390/math10214001.

[35] Shi Yuxin, Jiang Yue, Jiang Yue, et al. The optimal ratio of ginseng and astragalus to immune activity selected based on the entropy weight method and grey correlation degree method[J]. Chinese Journal of Hospital Pharmacy, 2025, 45(7): 738–745. DOI: 10.13286/j.1001-5213.2025.07.02.

[36] Li Z, Lin Y, Zou Y, et al. Zuogui pills ameliorate chemotherapy-induced ovarian aging by improving stemness, regulating cell cycle and reducing apoptosis of oogonial stem cells via the Notch1/Nrf2 pathway[J]. Journal of Ethnopharmacology, 2025, 339: 119105. DOI: 10.1016/j.jep.2024.119105.

[37] Ajonuma L C, Bamiro S A, Makanjuola S L. Prolonged use of pausinystalia yohimbe affects estrus cycle, reproductive hormones and folliculogenesis in rats[J]. Fertility and Sterility, 2018, 110(4): e381. DOI: 10.1016/j.fertnstert.2018.07.1067.

[38] Ma Q, Shen M, Wu J, et al. Mechanism research of DHEA treatment improving diminished ovarian reserve by attenuating the AMPK-SIRT1 signaling and mitophagy[J]. Reproductive Sciences, 2024, 31(7): 2059–2072. DOI: 10.1007/s43032-024-01473-3.

[39] Song Y, Hu R, Li F, et al. In view of ovarian steroidogenesis and luteal construction to explore the effects of bushen huoxue recipe in mice of ovarian hyperstimulation[J]. Journal of Ethnopharmacology, 2024, 318: 116913. DOI: 10.1016/j.jep.2023.116913.

[40] Chang H M, Klausen C, Leung P C K. Antimüllerian hormone inhibits follicle-stimulating hormone-induced adenylyl cyclase activation, aromatase expression, and estradiol production in human granulosa-lutein cells[J]. Fertility and Sterility, 2013, 100(2): 585–592.e1. DOI: 10.1016/j.fertnstert.2013.04.019.

[41] Nisa I C, Chantachotikul P, Takahata K, et al. SERPINE1 (PAI-1) regulation in mechanotransduction-associated cellular senescence[J]. Journal of Cellular Biochemistry, 2026, 127(9): e70123. DOI: 10.1002/jcb.70123.

[42] Tatone C, Amicarelli F. The aging ovary--the poor granulosa cells[J]. Fertility and Sterility, 2013, 99(1): 12–17. DOI: 10.1016/j.fertnstert.2012.11.029.

[43] Kim S, Pyun J A, Cha D H, et al. Epistasis between FSHR and CYP19A1 polymorphisms is associated with premature ovarian failure[J]. Fertility and Sterility, 2011, 95(8): 2585–2588. DOI: 10.1016/j.fertnstert.2010.12.042.

[44] Mohyeldin R H, Sharata E E, Abdelnaser M, et al. Imeglimin halts cyclophosphamide-induced ovarian injury: a dual modulation of TLR4/NF-κB/NLRP3-driven inflammation and AMPK/SIRT1/PGC-1α pathways[J]. Naunyn-Schmiedeberg’s Archives of Pharmacology, 2026. DOI: 10.1007/s00210-026-05885-7.

[45] Blengini C S, Schindler K. Follicular communication breakdown in aging ovaries[J/OL]. Nature Aging, 2023, 3(6): 636–637. DOI: 10.1038/s43587-023-00435-9.

[46] Ju W, Zhao Y, Yu Y, et al. Mechanisms of mitochondrial dysfunction in ovarian aging and potential interventions[J]. Frontiers in Endocrinology, 2024, 15: 1361289. DOI: 10.3389/fendo.2024.1361289.

[47] Siyu Y, Shixiao Z, Congying S, et al. Advances in cytokine-based herbal medicine against premature ovarian insufficiency: a review[J]. Journal of Ethnopharmacology, 2024, 333: 118477. DOI: 10.1016/j.jep.2024.118477.

[48] Song Z, Chen G, Chen C Y C. AI empowering traditional chinese medicine?[J]. Chemical Science, 2024, 15(41): 16844–16886. DOI: 10.1039/d4sc04107k.

[49] Navarro-Pando J M, Alcocer-Gómez E, Castejón-Vega B, et al. Inhibition of the NLRP3 inflammasome prevents ovarian aging[J]. Science Advances, 2021, 7(1): eabc7409. DOI: 10.1126/sciadv.abc7409.

[50] Yunhui L, Zhou X, Ma L, et al. O-078 deletion of interlukin-17A protects against ovarian aging via activating wnt10b/β-catenin signaling[J]. Human Reproduction, 2024, 39(Supplement_1): deae108.084. DOI: 10.1093/humrep/deae108.084.

[51] Markova I, Hüttl M, Gayova N, et al. Visceral adipose tissue inflammation and vascular complications in a rat model with severe dyslipidemia: Sex differences and PAI-1 tissue involvement[J]. Biomolecules, 2024, 15(1): 19. DOI: 10.3390/biom15010019.

[52] Simone T, Higgins P. Inhibition of SERPINE1 function attenuates wound closure in response to tissue injury: A role for PAI-1 in re-epithelialization and granulation tissue formation[J]. Journal of Developmental Biology, 2015, 3(1): 11–24. DOI: 10.3390/jdb3010011.

[53] Khan S S, Shah S J, Klyachko E, et al. A null mutation in *SERPINE1* protects against biological aging in humans[J]. Science Advances, 2017, 3(11): 1617. DOI: 10.1126/sciadv.aao1617.

[54] Wu J, Wang J, Pei Z, et al. Endothelial senescence induced by PAI-1 promotes endometrial fibrosis[J]. Cell Death Discovery, 2025, 11(1): 89. DOI: 10.1038/s41420-025-02377-0.

[55] Zhang K H, Zhang F F, Zhang Z L, et al. Follicle stimulating hormone controls granulosa cell glutamine synthesis to regulate ovulation[J]. Protein & Cell, 2024, 15(7): 512–529. DOI: 10.1093/procel/pwad065.

[56] Yu X, Xie L, Ge J, et al. Integrating single-cell RNA-seq and spatial transcriptomics reveals MDK-NCL dependent immunosuppressive environment in endometrial carcinoma[J]. Frontiers in Immunology, 2023, 14: 1145300. DOI: 10.3389/fimmu.2023.1145300.

[57] Fei H, Chen T, Jiang H. Autocrine and paracrine effects of MDK promote lymph node metastasis of cervical squamous cell carcinoma[J]. iScience, 2024, 27(7): 110077. DOI: 10.1016/j.isci.2024.110077.

[58] Liu X, Onda M, Watson N, et al. Highly active CAR T cells that bind to a juxtamembrane region of mesothelin and are not blocked by shed mesothelin[J]. Proceedings of the National Academy of Sciences of the United States of America, 2022, 119(19): e2202439119. DOI: 10.1073/pnas.2202439119.

[59] János L. Tanyi, Andrew R. Haas, Charu Aggarwal, et al. Phase I study of autologous T cells bearing fully-humanized chimeric antigen receptors targeting mesothelin in mesothelin-expressing cancers (314)[J]. GYNECOLOGIC ONCOLOGY. 2022, 166(0): S164–S165. DOI: 10.1016/s0090-8258(22)01537-2.

[60] Hilliard T S. The impact of mesothelin in the ovarian cancer tumor microenvironment[J]. Cancers, 2018, 10(9): 277. DOI: 10.3390/cancers10090277.

[61] Weckbach L T, Gola A, Winkelmann M, et al. The cytokine midkine supports neutrophil trafficking during acute inflammation by promoting adhesion via β2 integrins (CD11/CD18)[J]. Blood, 2014, 123(12): 1887–1896. DOI: 10.1182/blood-2013-06-510875.

[62] Al-Zoubi A M, Mora A M, Faris H, et al. A hybrid TwinSVM-HHO model for multilingual spam review detection using sentiment features and pre-trained embeddings[J]. Expert Systems with Applications, 2025, 287: 128160. DOI: 10.1016/j.eswa.2025.128160.

[63] Gao W. Comprehensive summary of material basis – laying a foundation for substances of medicine food homology development[J]. Chinese Herbal Medicines, 2024, 16(3): 311–312. DOI: 10.1016/j.chmed.2024.05.001.

[64] Chen J. Essential role of medicine and food homology in health and wellness[J]. Chinese Herbal Medicines, 2023, 15(3): 347–348. DOI: 10.1016/j.chmed.2023.05.001.

[65] Zhu L, Luo S P. Professor Luo Songping’s Treatment of Premature Ovarian Failure Based on Yin and Yang[J]. Journal of Basic Chinese Medicine, 2020, 26(6): 841–843.

[66] Xu Z L, Wang Y. Treating Premature Ovarian Failure from the Perspective of Spleen [J]. Inner Mongolia Journal of Traditional Chinese Medicine, 2024, 43(9): 148–151. DOI: 10.16040/j.cnki.cn15-1101.2024.09.014.

[67] Han Y H, Geng T T, Han Y G. HANs’ gynecology experience in treating premature ovarian failure based on the theory of ‘liver and kidney theory’ [J]. China Journal of Traditional Chinese Medicine and Pharmacy, 2021, 36(12): 7177–7179.

[68] Li A, Wang X, Yang R, et al. Therapeutic potential and mechanisms of traditional chinese medicine in regulating energy metabolism imbalance in heart failure[J]. Chinese Herbal Medicines, 2025, 17(4): 685–702. DOI: 10.1016/j.chmed.2025.07.002.

[69] Zhang X, Lin B, Wang X, et al. Research Progress on the Treatment of Related Diseases With Astragalus[J]. Drug Design, Development and Therapy, 2025, 19: 2845–2862. DOI: 10.2147/dddt.s494915.

[70] Tian M, Li L N, Zheng R R, et al. Advances on hormone-like activity of Panax ginseng and ginsenosides[J]. Chinese Journal of Natural Medicines, 2020, 18(7): 526–535. DOI: 10.1016/s1875-5364(20)30063-7.

[71] Zhang T, An W, You S, et al. G protein-coupled receptors and traditional chinese medicine: new thinks for the development of traditional chinese medicine[J]. Chinese Medicine, 2024, 19(1): 92. DOI: 10.1186/s13020-024-00964-4.

[72] Chen Y, Wang S, Zhang C. The differentiation fate of granulosa cells and the regulatory mechanism in ovary[J]. Reproductive Sciences, 2025, 32(5): 1414–1426. DOI: 10.1007/s43032-024-01682-w.

[73] Zhang Y, Yan Z, Qin Q, et al. Transcriptome landscape of human folliculogenesis reveals oocyte and granulosa cell interactions[J/OL]. Molecular Cell, 2018, 72(6): 1021–1034. DOI: 10.1016/j.molcel.2018.10.029.

[74] Hirano M, Onodera T, Takasaki K, et al. Ovarian aging: Pathophysiology and recent developments in maintaining ovarian reserve[J]. Frontiers in Endocrinology, 2025, 16: 1619516. DOI: 10.3389/fendo.2025.1619516.

[75] Isola J V V, Hense J D, Osório C A P, et al. Reproductive ageing: Inflammation, immune cells, and cellular senescence in the aging ovary[J]. Reproduction, 2024, 168(2): e230499. DOI: 10.1530/REP-23-0499.

[76] Sharata E E, Attya M E, Khalaf M M, et al. Decoding premature ovarian insufficiency: A full spectrum review from epidemiology to management with emphasis on cyclophosphamide-evoked ovarian injury[J]. Food and Chemical Toxicology, 2025, 205: 115719. DOI: 10.1016/j.fct.2025.115719.

[77] Sharata E E, Bakry T, Atta H G, et al. Molecular mechanisms underlying cyclophosphamide-induced ovarian injury and protective strategies[J]. Naunyn-Schmiedeberg’s Archives of Pharmacology, 2026, 399(2): 1951–1985. DOI: 10.1007/s00210-025-04550-9.

