## Supplementary Materials for "Screening and evaluation of medicine-food homology herbal combinations for ameliorating chemotherapy-induced ovarian aging under an integrated multi-strategy framework"

**Supplementary Figures**

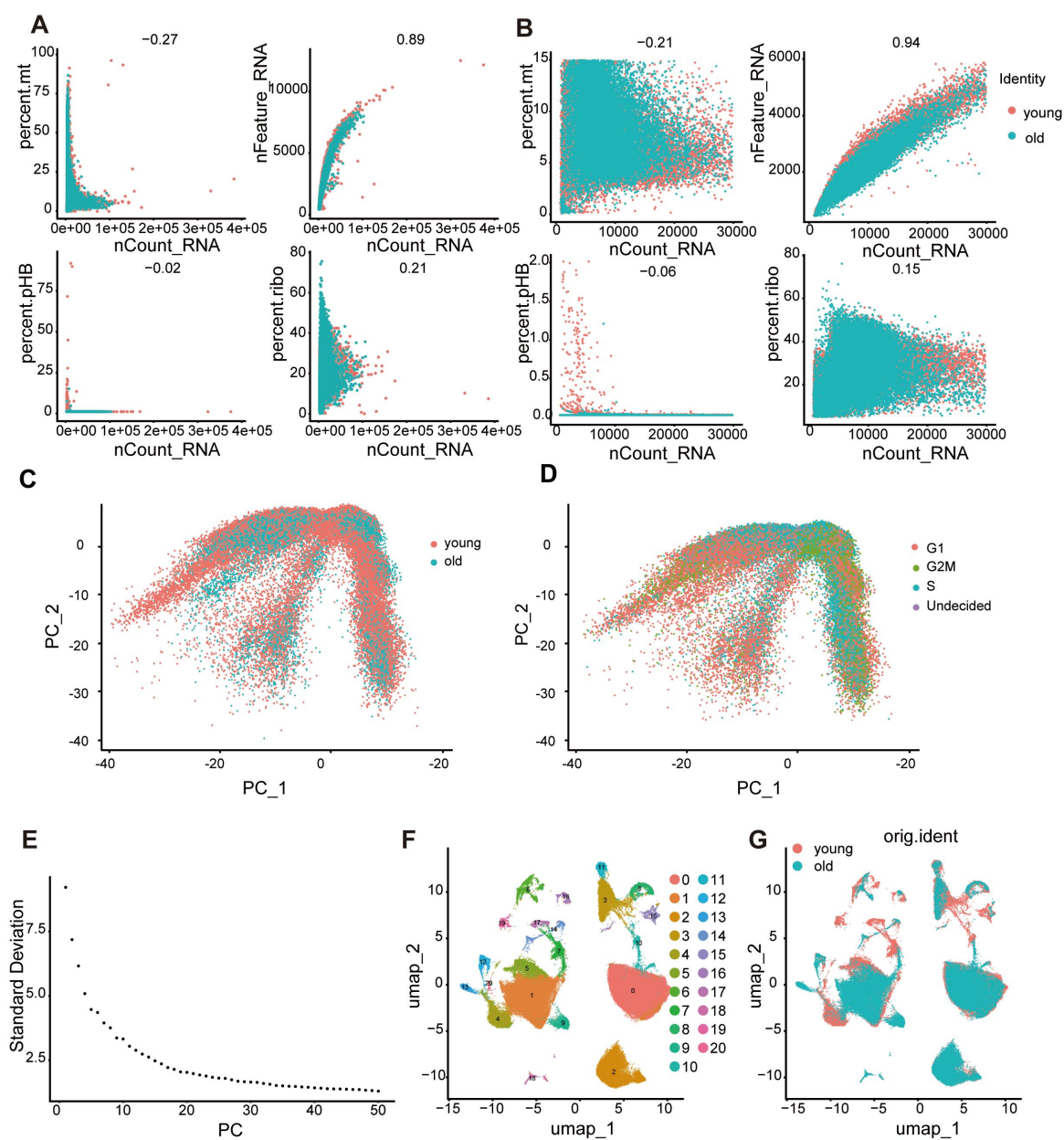

**Supplementary Figure 1. Quality control, batch effect assessment and dimensionality reduction of ovarian scRNA-seq data.** (A, B) Scatter plots showing the correlation of mitochondrial RNA percentage (percent.mt), detected gene number (nFeature\_RNA), hemoglobin RNA percentage (percent.pHB) and ribosomal RNA percentage (percent.ribo) with UMI counts (nCount\_RNA) before (A) and after (B) quality control, with correlation coefficients shown at the top of each panel. (C, D) PCA plots colored by sample group (C) and cell-cycle phase (D), showing no evident separation by batch or cell-cycle stage after

integration. (E) Scree plot of standard deviation for the first 50 principal components; the
top 1–50 PCs were selected for downstream clustering. (F) UMAP plot of the 20 identified
cell clusters. (G) UMAP plots of young and old groups, showing well-overlapped cell
distributions after integration.

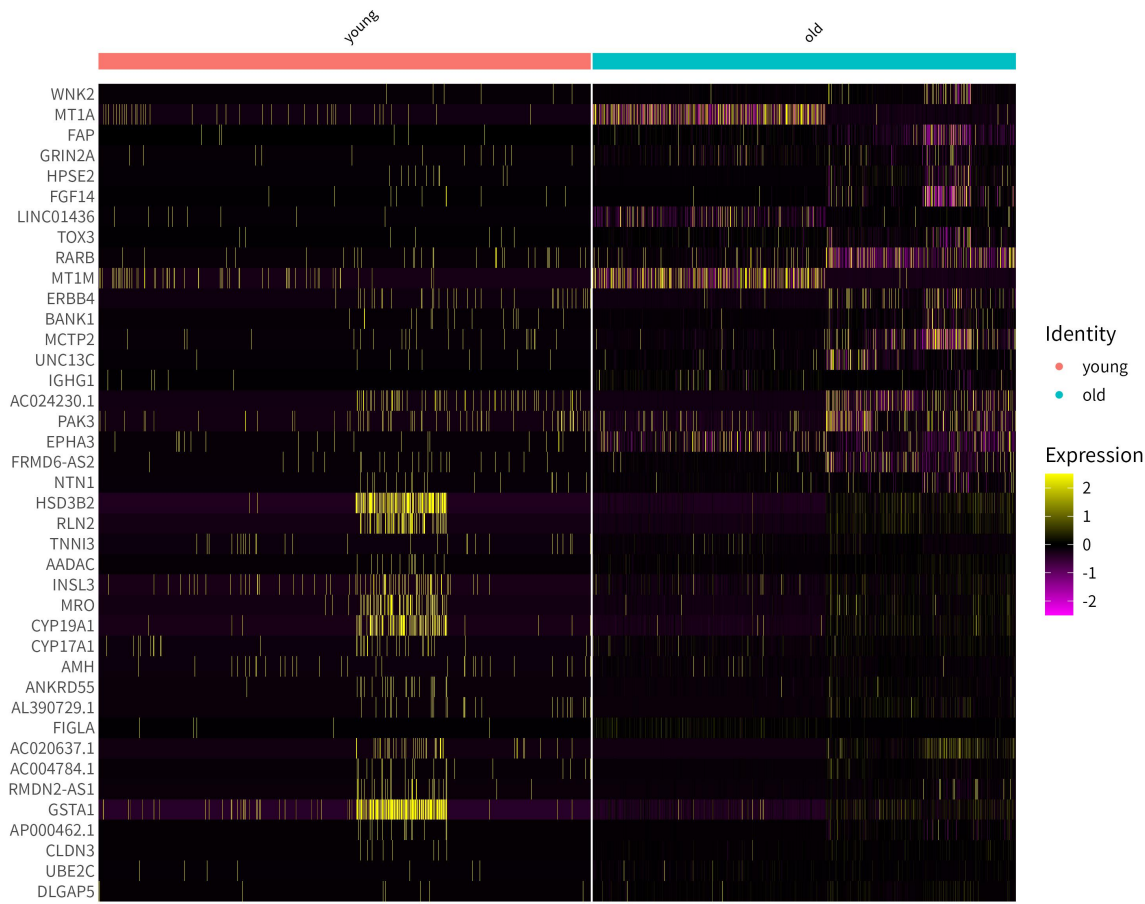

**Supplementary Figure 2. Global differential gene analysis (DEGs)**

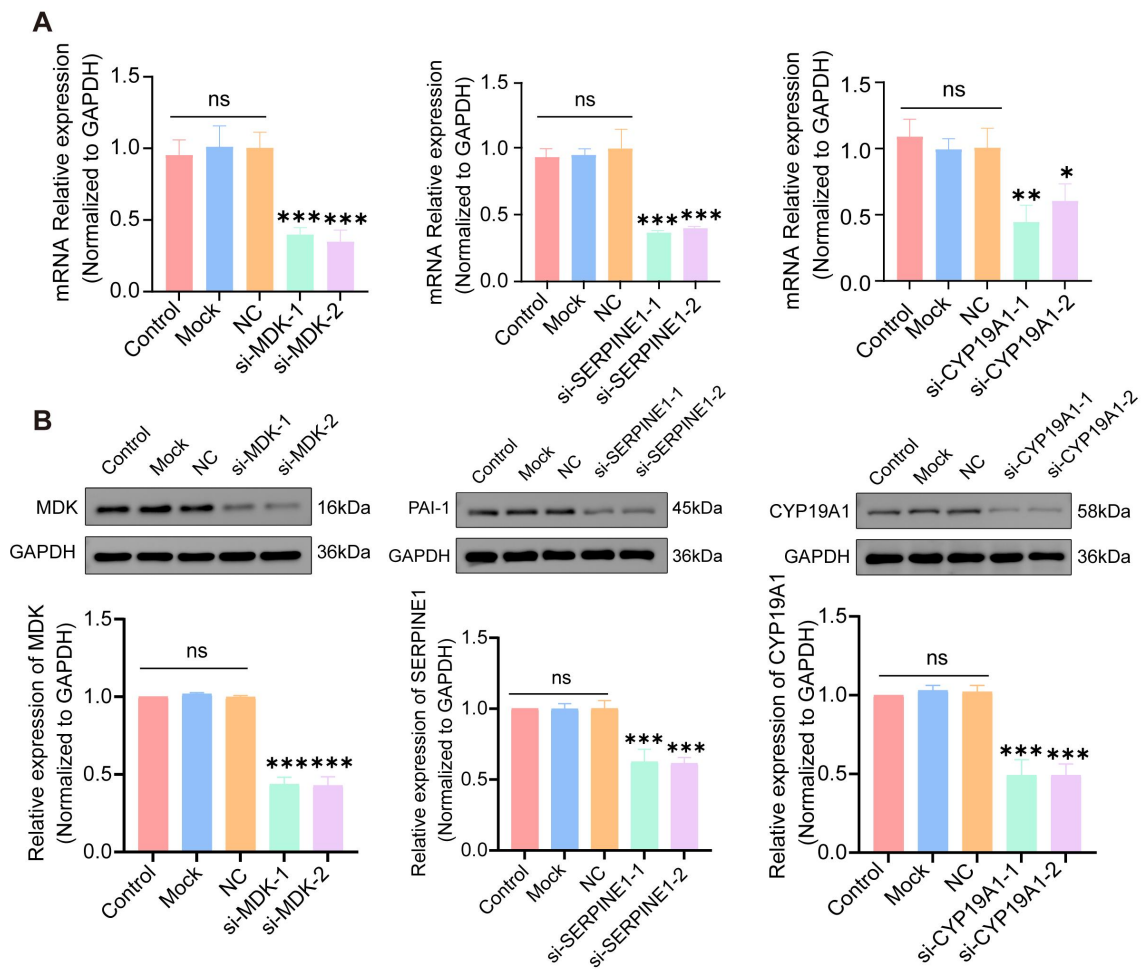

**Supplementary Figure 3. Knockdown efficiency of siRNAs targeting MDK, SERPINE1 and CYP19A1 in KGN cells. (A) Relative mRNA expression of MDK, SERPINE1 and CYP19A1 detected by qRT-PCR. (B) Western blot results corresponding to (A), showing MDK, SERPINE1 and CYP19A1 protein levels in each group.**

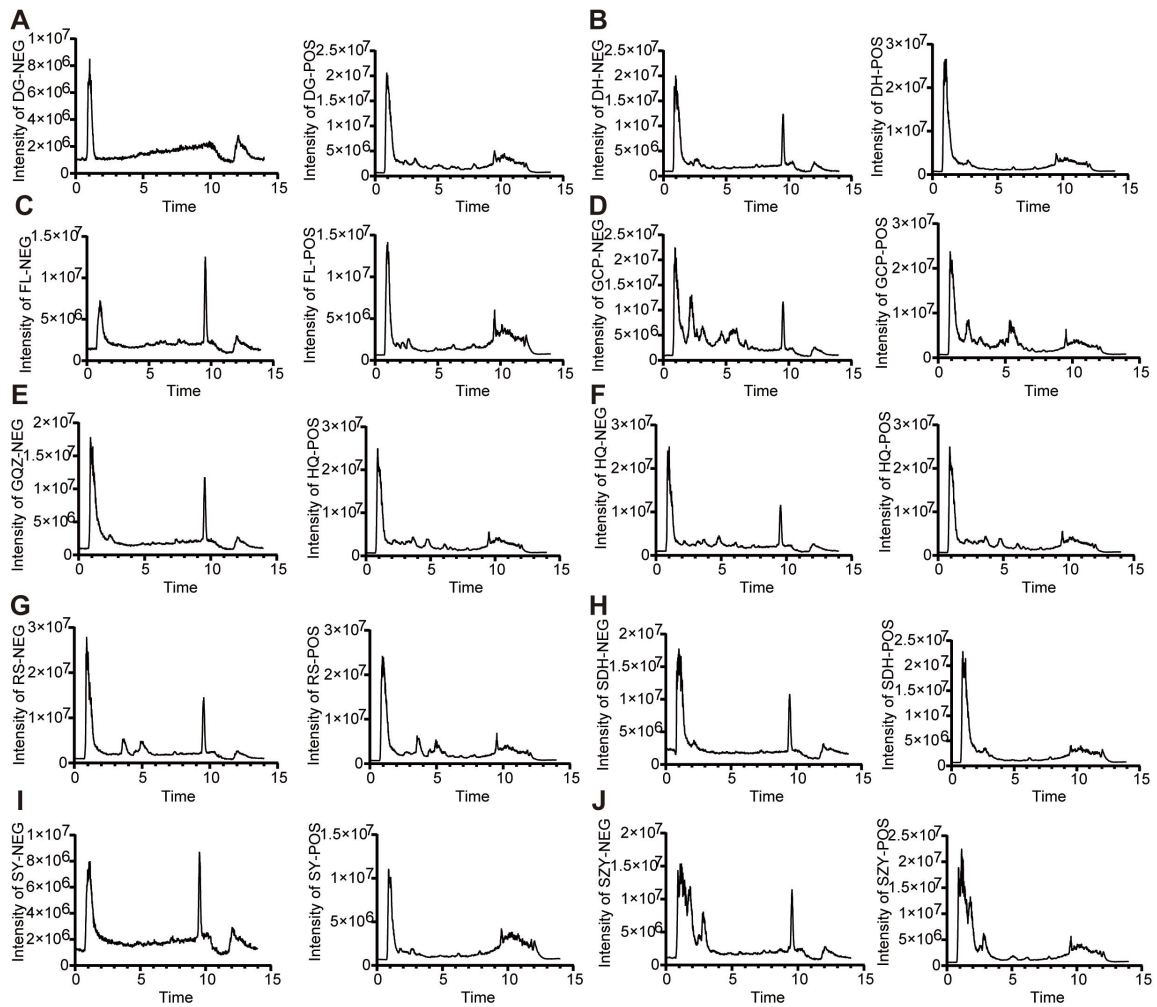

**Supplementary Figure 4. Total ion chromatogram of decoction from ten Chinese medicinal herbs.** (A) Angelicae Sinensis Radix(DG); (B) Rehmanniae Radix(DH); (C) Poria(FL); (D) Ginseng Radix et Rhizoma(GCP); (E) Lyci Fructus(GQZ); (F) Astragali Radix(HQ); (G) Glycyrrhizae Radix et Rhizoma(RS); (H) Rehmanniae Radix Praeparata(SDH); (I) Dioscoreae Rhizoma(SY); (J) Corni Fructus(SZY).

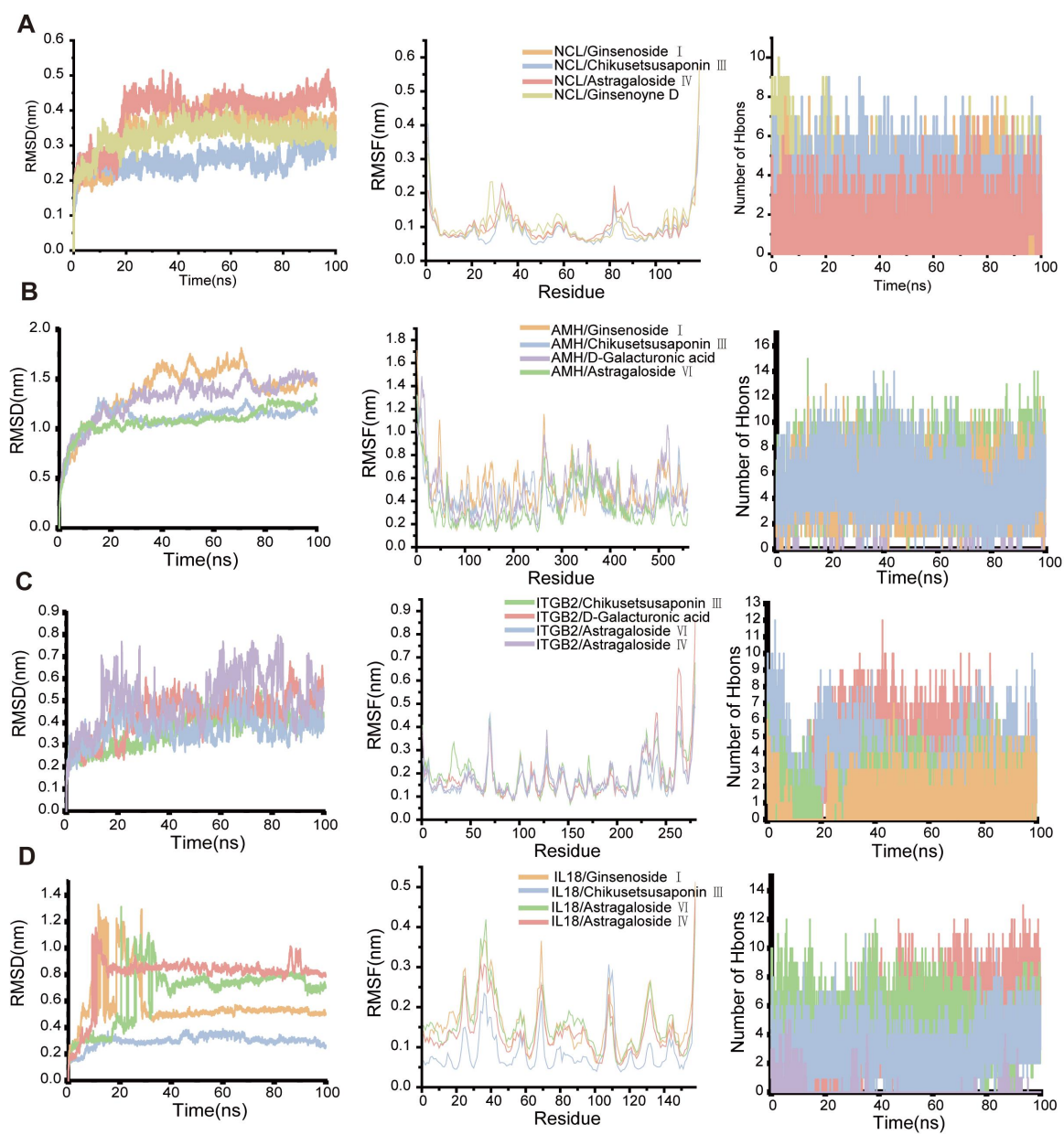

**Supplementary Figure 5. Molecular dynamics simulation analysis of key active components from HXF with target proteins. (A–D) Protein–ligand complexes of NCL, AMH, ITGB2, and IL1B with representative active components, respectively. From left to right in each row: root mean square deviation (RMSD), residue root mean square fluctuation (RMSF), and hydrogen bond number changes during 100 ns molecular dynamics simulations.**

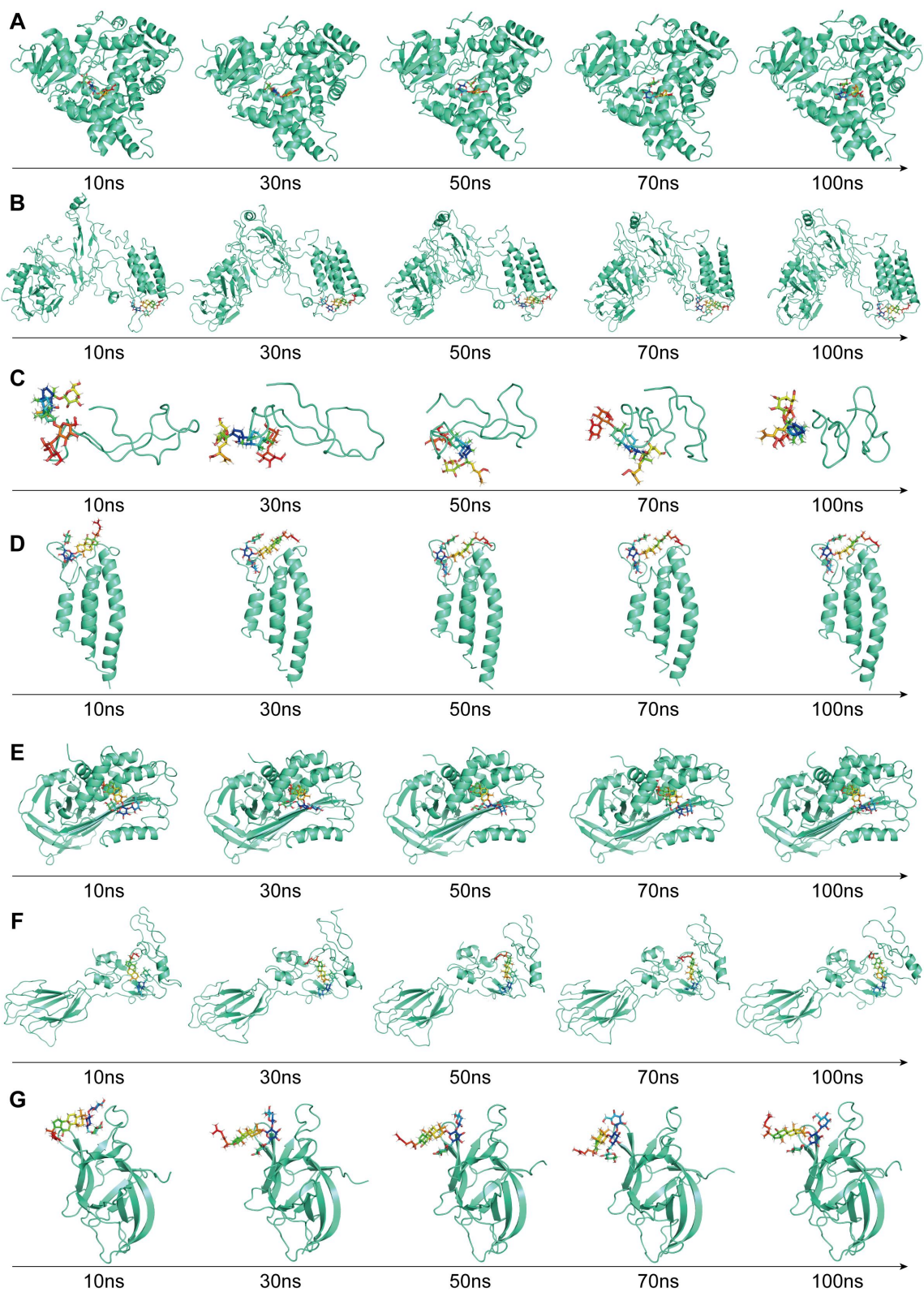

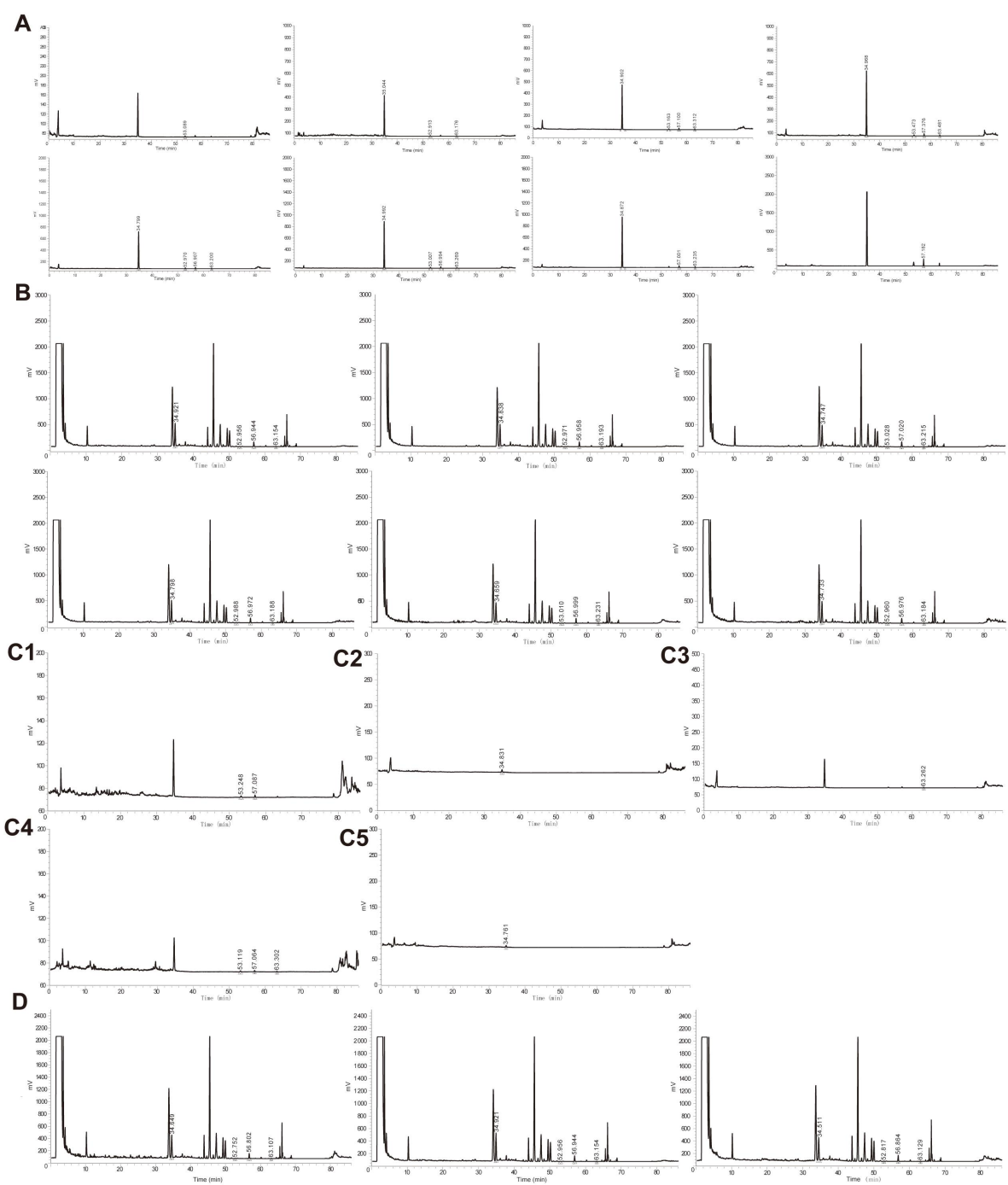

**Supplementary Figure 7. Chromatograms of the HPLC-ELSD method validation for the simultaneous quantification of ginsenoside Re, astragaloside IV, ginsenoside Rd, and isoastragaloside II in HXF.** (A) HPLC chromatograms of mixed standard solutions at a series of concentrations used for establishing the calibration curves ($R^2 \geq 0.99$ for all four analytes). (B) Chromatograms of six replicate injections of the HXF2 sample solution (precision test; the RSDs of the contents of the four analytes were all below 2%). (C) Chromatograms of the limits of quantification (LOQ) and limits of detection (LOD) of each analyte: (C1) LOQ of astragaloside IV (0.187 $\mu\text{g}$) and ginsenoside Rd (0.165 $\mu\text{g}$); (C2) LOQ of ginsenoside Re (0.396 $\mu\text{g}$); (C3) LOQ of isoastragaloside II (0.138 $\mu\text{g}$); (C4) LOD of astragaloside IV, ginsenoside Rd, and isoastragaloside II (0.117, 0.103, and 0.0574 $\mu\text{g}$, respectively); (C5) LOD of ginsenoside Re (0.297 $\mu\text{g}$). (D) Representative chromatograms of the three batches of HXF samples (HXF1–HXF3) for content determination.

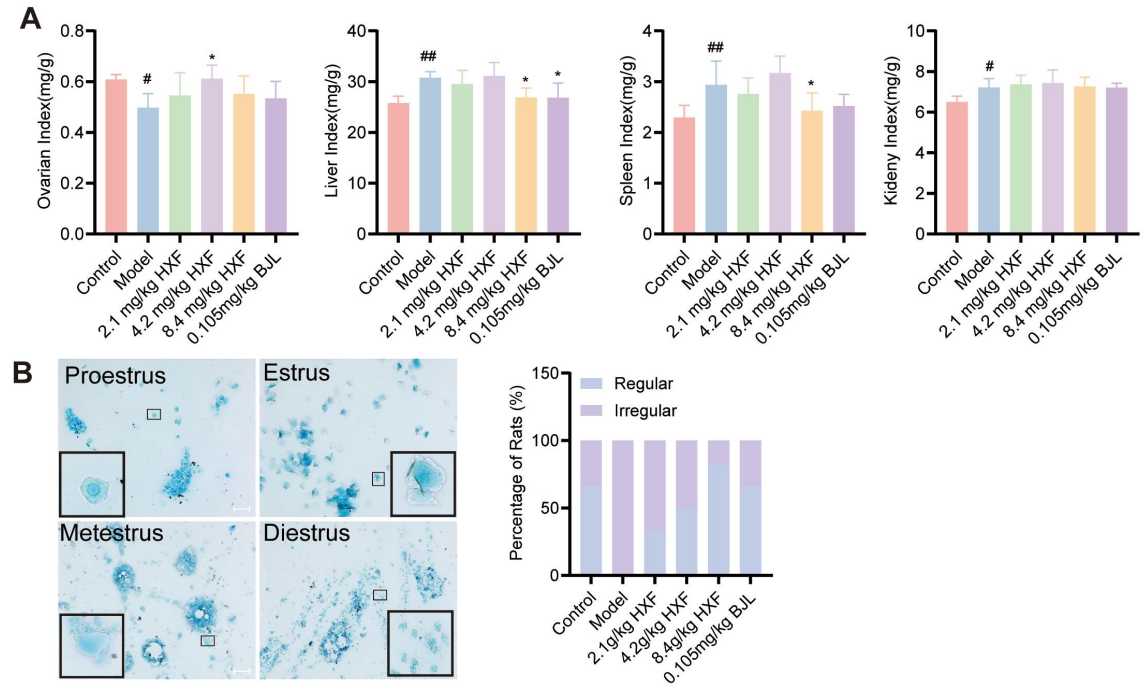

**Supplementary Figure 8. Effects of HXF on organ indexes and estrous cyclicity in model rats.** (A) Ovarian, liver, spleen, and kidney indexes of rats in each group ($n = 6$). (B) Representative vaginal smear cytology showing the typical cellular patterns of each estrous stage, and the percentage of rats with regular or irregular estrous cycles in each group. #$P < 0.05$, ##$P < 0.01$ vs. the Control group; \*$P < 0.05$ vs. the Model group.

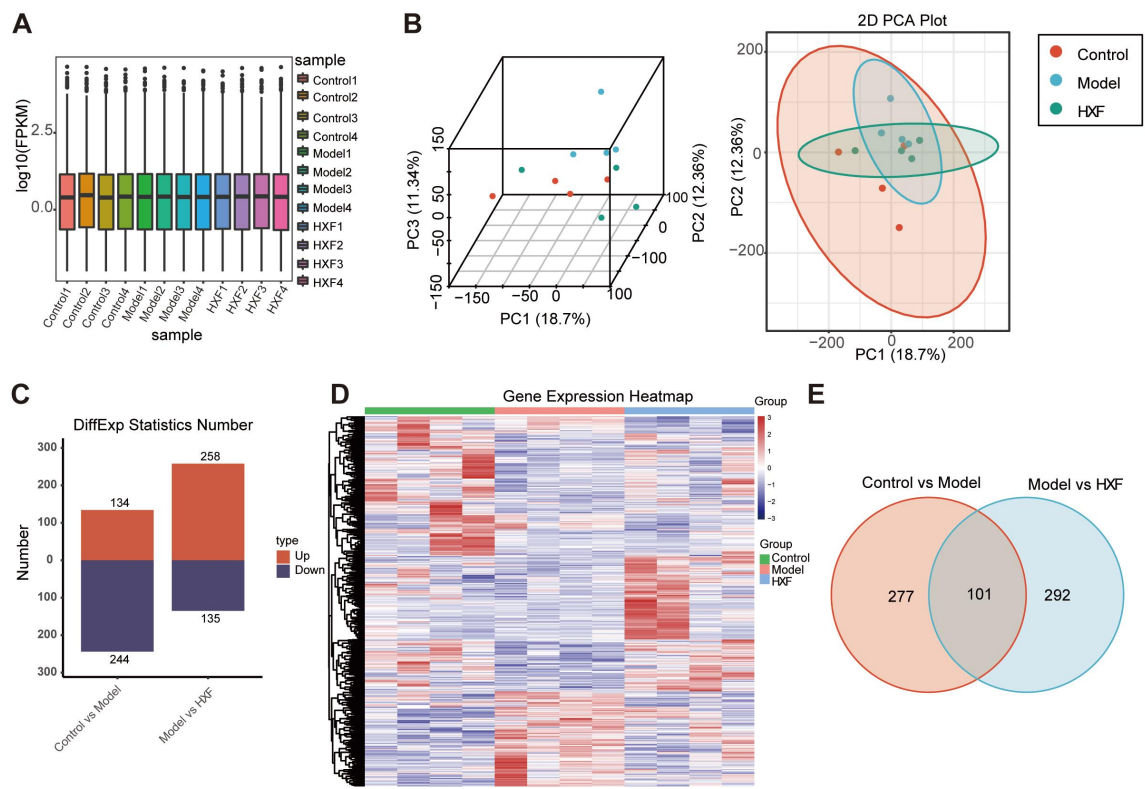

**Supplementary Figure 9. Quality assessment of the rat ovarian transcriptome sequencing data and analysis of differentially expressed genes (DEGs).** (A) Box plots of the $\log_{10}(\text{FPKM})$ distribution for each sample (n = 4 per group). (B) Three-dimensional and two-dimensional principal component analysis (PCA) plots of the samples. (C) Numbers of upregulated and downregulated genes. (D) Heatmap of DEGs. (E) Venn diagram of DEGs identified in the two comparisons.

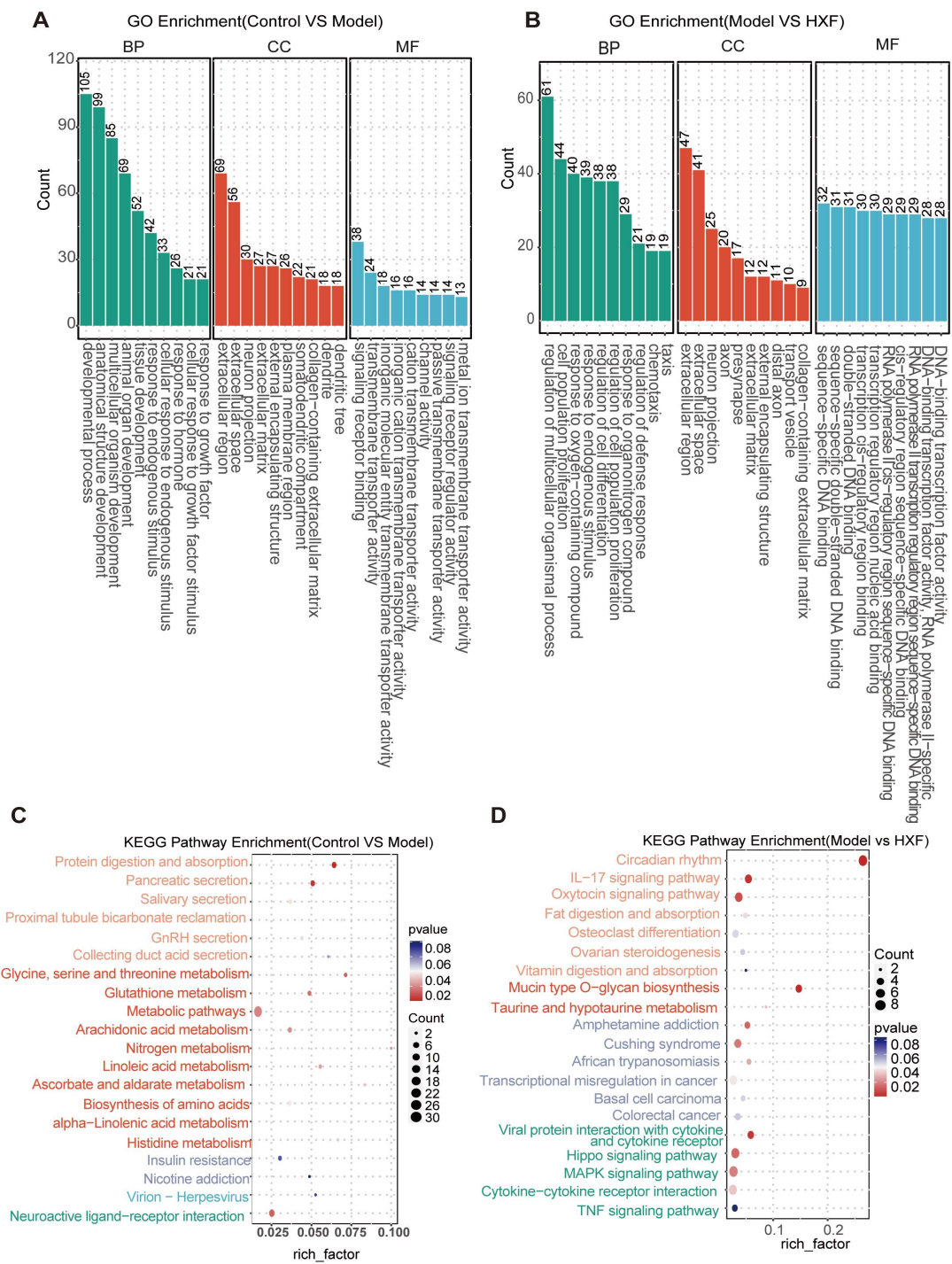

**Supplementary Figure 10. GO functional enrichment and KEGG pathway analyses of the differentially expressed genes.** (A) GO enrichment analysis of DEGs in the Control vs. Model comparison. (B) GO enrichment analysis of DEGs in the Model vs. HXF comparison. In (A, B), the top 10 enriched terms are presented for biological process (BP),

cellular component (CC), and molecular function (MF). (C) KEGG pathway enrichment
bubble plot of DEGs in the Control vs. Model comparison. (D) KEGG pathway enrichment
bubble plot of DEGs in the Model vs. HXF comparison. Bubble size indicates the number
of enriched genes, and bubble color indicates the P value.

**Supplementary Table 1. Major constituents in the aqueous extracts of each medicinal material and their relative retention times ( part )**

| 序号 | Smiles_id | Name | Drug | Formula | Adduct | Extraction Mass(Da) | Found At Mass(Da) | $\delta$/ppm | RT/min |
| --- | --- | --- | --- | --- | --- | --- | --- | --- | --- |
| 1 | ID0053 | 3,4,6-trihydroxyphenanthrene-3-o-<br>β-d-glucopyranoside | SY | C<sub>20</sub>H<sub>20</sub>O<sub>8</sub> | [M-H]- | 387.10854 | 387.1133 | 12.3 | 0.91 |
| 2 | ID0059 | Allantoin | SY | C<sub>4</sub>H<sub>6</sub>N<sub>4</sub>O<sub>3</sub> | [M-H]- | 157.03671 | 157.03756 | 5.4 | 0.98 |
| 3 | ID0009 | L-Xyl | SY | C<sub>5</sub>H<sub>10</sub>O<sub>5</sub> | [M-H]- | 149.04555 | 149.04644 | 6 | 1.02 |
| 4 | ID0027 | Batatasin IV | SY | C<sub>15</sub>H<sub>16</sub>O<sub>3</sub> | [M-H]- | 243.10267 | 243.10277 | 0.4 | 4.7 |
| 5 | ID0029 | Batatasin I | SY | C<sub>6</sub>H<sub>14</sub>N<sub>2</sub>O<sub>2</sub> | [M+H]+ | 147.1128 | 147.11273 | -0.5 | 0.87 |
| 6 | ID0006 | Leucinum | SY | C<sub>10</sub>H<sub>16</sub>O<sub>3</sub> | [M+H]+ | 185.11722 | 185.11521 | -10.9 | 3.15 |
| 7 | ID0008 | L-Valin | SY | C<sub>6</sub>H<sub>13</sub>NO<sub>2</sub> | [M+H]+ | 132.10191 | 132.10181 | -0.7 | 1.14 |
| 8 | ID0007 | L-Lysin | SY | C<sub>5</sub>H<sub>11</sub>NO<sub>2</sub> | [M+H]+ | 118.08626 | 118.08616 | -0.8 | 1.1 |
| 9 | ID0001 | BGC | SY | C<sub>6</sub>H<sub>12</sub>O<sub>6</sub> | [M-H]- | 179.05611 | 179.05711 | 5.6 | 0.95 |
| 10 | ID0063 | Batatasin v | SY | C<sub>17</sub>H<sub>20</sub>O<sub>4</sub> | [M+H]+ | 289.14344 | 289.14249 | -3.3 | 5.48 |
| 11 | ID1242 | Raffinose | SDH | C<sub>18</sub>H<sub>32</sub>O<sub>16</sub> | [M-H]- | 503.16176 | 503.1622 | 0.9 | 0.88 |
| 12 | ID1124 | Sucrose | SDH | C<sub>12</sub>H<sub>22</sub>O<sub>11</sub> | [M-H]- | 341.10894 | 341.10885 | -0.2 | 0.92 |

**Supplementary Table 2. Predicted pKD values for selected target–compound pairs (part)**

| Protein | Smiles_id | Name | Predicted_pKD | Drug from |
| --- | --- | --- | --- | --- |
| PAI-1 | ID1225 | Chikusetsusaponin III | 10.093496 | RS |
|  | ID1326 | Astragaloside VI | 9.858257 | HQ |
|  | ID1374 | Astragaloside IV | 9.793938 | HQ |
|  | ID1318 | Astragaloside V | 9.730873 | HQ |
|  | ID1621 | Rehmannioside D | 9.586362 | SDH |
|  | ID1118 | Ginsenoside I | 9.555254 | RS |
|  | ID1308 | Astragaloside III | 9.466071 | HQ |
|  | ID0152 | Ginsenoside Rd | 9.3168125 | RS |
|  | ID1140 | Notoginsenoside R6 | 9.274109 | RS |
|  | ID1577 | Cistanoside A | 9.244301 | SDH |
|  | ID1225 | Chikusetsusaponin III | 9.989513 | RS |
| NCL | ID1374 | Astragaloside IV | 9.752062 | HQ |
|  | ID1326 | Astragaloside VI | 9.533809 | HQ |
|  | ID1118 | Ginsenoside I | 9.418791 | RS |
|  | ID1318 | Astragaloside V | 9.371032 | HQ |
|  | ID0153 | Ginsenoside Re | 9.271929 | RS |
|  | ID1136 | Ginsenoside Ic | 9.255922 | RS |
|  | ID0152 | Ginsenoside Rd | 9.204796 | RS |
|  | ID1621 | Rehmannioside D | 9.150141 | SDH |
|  | ID1174 | Pseudoginsenoside F11 | 9.079836 | RS |
|  | ID1199 | Nonacosanediol-6,8 | 9.129414 | RS |
| MDK | ID1225 | Chikusetsusaponin III | 8.772648 | RS |
|  | ID0291 | 1-Acetyl-4,6,8-Trimethylazulene | 8.742983 | SZY |
